# Deciphering *Legionella* effector delivery by Icm/Dot secretion system reveals a new role for c-diGMP signaling

**DOI:** 10.1101/754762

**Authors:** J. Allombert, C. Jaboulay, C. Michard, C. Andréa, X. Charpentier, A. Vianney, P. Doublet

## Abstract

Secretion of bacterial effector proteins into host cells plays a key role in bacterial virulence. Yet, the dynamics of the secretion systems activity remains poorly understood, especially when machineries deal with the export of numerous effectors. We address the question of multi-effector secretion by focusing on the *Legionella pneumophila* Icm/Dot T4SS that translocates a record number of 300 effectors. We set up a kinetic translocation assay, based on the *β*-lactamase translocation reporter system combined with the effect of the protonophore CCCP. When used for translocation analysis of Icm/Dot substrates constitutively produced by *L. pneumophila,* this assay allows a fine monitoring of the secretion activity of the T4SS, independently of the expression control of the effectors. We observed that effectors are translocated with a specific timing, suggesting a control of their docking/translocation by the T4SS. Their delivery is accurately organized to allow effective manipulation of the host cell, as exemplified by the sequential translocation of effectors targeting Rab1, namely SidM/DrrA, LidA, LepB. Remarkably, the timed delivery of effectors does not depend only on their interaction with chaperone proteins but implies cyclic-di-GMP signaling, as the diguanylate cyclase Lpl0780/Lpp0809, contributes to the timing of translocation.

**Graphical abstract:** 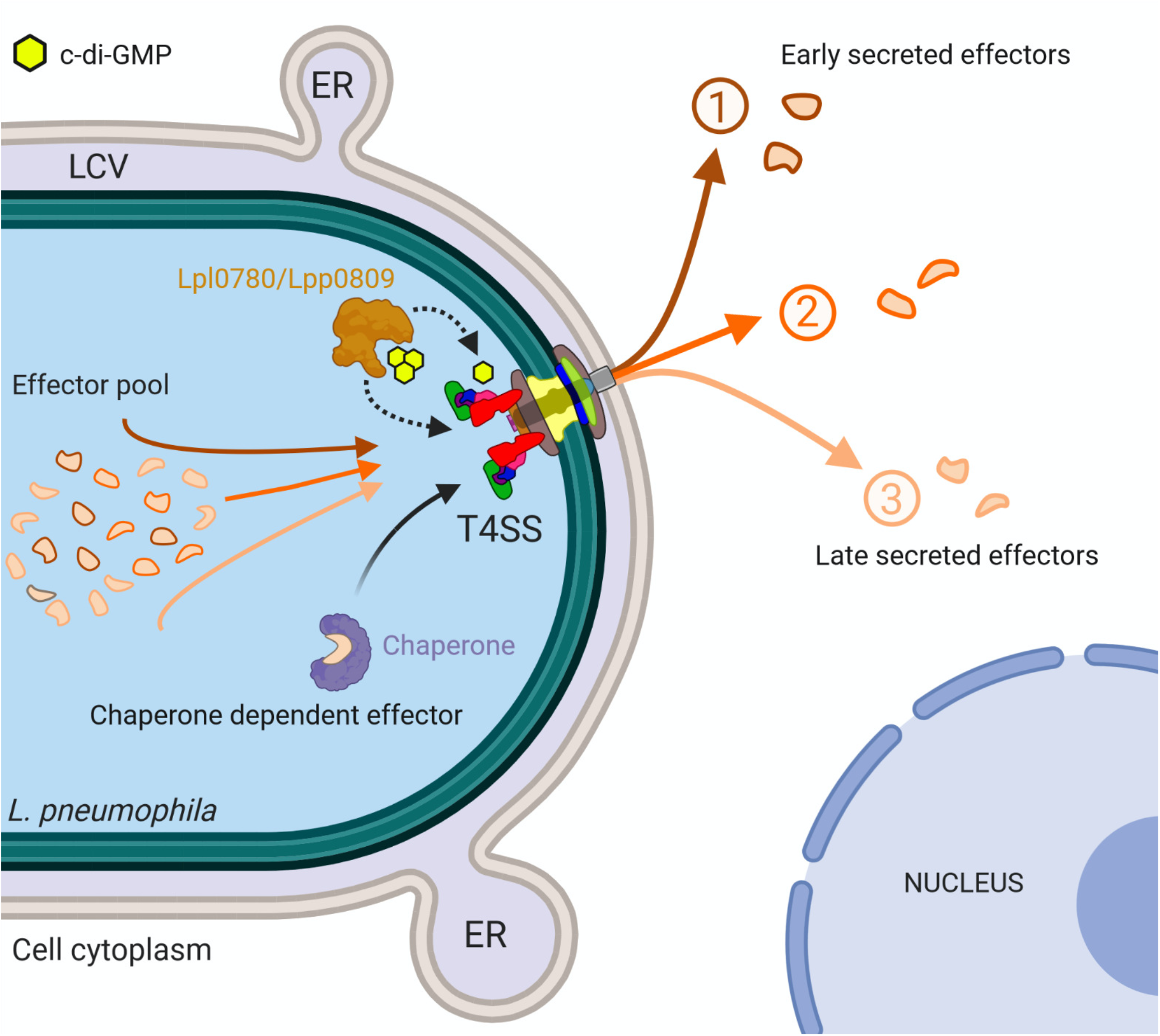

**Highlights:**

- The dynamics of multi-effector secretion is addressed in the paradigm *Legionella* Icm/Dot T4SS
- A kinetic translocation assay allowing a fine monitoring of the T4SS activity is set up
- Specific secretion patterns consistent with sequential functions in the host are reported
- T4SS-dependent translocation is promoted by a diguanylate cyclase
- Unreported control of Type 4 secretion by the second messenger c-di-GMP signaling is revealed

## Introduction

Delivery of effector proteins that hijack host cell processes to the benefit of the bacteria is a mechanism widely used by bacterial pathogens. Effector delivery is achieved by complex effector injection devices such as the Type III, the Type IV and the Type VI secretion systems (T3SS, T4SS, T6SS). Although considerable progresses have been made in structure-function analysis of these secretion systems, few mechanisms leading to a sequential delivery of the effectors according to their role during infection are described. Most of them concern the T3SS. For example, *Salmonella* successively expresses two T3SS and their cognate effectors at the initial extracellular stage and later intracellular stage of infection [1]. In addition, T3SS sorting platform could allow sequential loading of the substrates [2]. In EPEC, the order of effectors translocation is governed by interactions with the CesT chaperone which ensure a priority delivery to the Tir effector by the T3SS [3]. Considering that these T3SS translocate fewer than a dozen of effectors, the case of *L. pneumophila* Icm/Dot T4SS is particularly intriguing as it injects more than 300 effectors in the host cell [4], a subset of which have been shown to be critical for hijacking host-cell vesicles trafficking necessary for *Legionella*-containing vacuole (LCV) biogenesis, and subsequently for intracellular multiplication [5, 6]. This complex machinery is composed of 25 proteins related to the multiprotein secretion apparatus [7] or the coupling protein complex (IcmO/DotL; IcmP/DotM; IcmJ/DotN; DotY; DotZ) [8–10]. Chaperone proteins (IcmS; IcmW; LvgA) that associate with the coupling protein complex are involved in some specific substrate recognition for presentation to the translocon [10–13]. Electron cryotomography has revealed its *in situ* polar molecular architecture with a central channel through which effectors would be transported to the host cytosol [14–18].

Based on the observation of the translocation of some effectors, it appears that effector translocation occurs rapidly upon phagocytosis [19]. Consistently, most of the Icm/Dot component encoding genes are continuously expressed during the intracellular cycle of the bacteria and the secretion machinery is thus assembled for the next cell infection [20]. Yet, it has been shown that not all effectors are simultaneously translocated into the host cytosol at the onset of infection and that a temporal control of the effector activity is required for *Legionella* to effectively manipulate the host cell pathways. The Icm/Dot system is active in effector translocation for at least 8 hours after bacterial uptake [21] and the effectors SidM/DrrA and SidE family members can be detected in proximity to the LCV early on but then disappear at later stages during infection, while LepB displays increased co-localization with the LCV over time [22–24]. Most importantly, the temporal control of the presence of some effectors in the host cell has been shown to be consistent with their biological function, as exemplified by the set of effectors SidM, LepB and SidD that target Rab1 with antagonist activities to temporally control its activation/desactivation on the LCV [23], or SidJ that inhibits the ubiquitin ligase activity of the SidE family effectors [25–28].

In addition to post-transcriptional regulation of effector level in the host cell by host proteasome or metaeffectors [29, 30], a temporal control of the effector activity might be primarily achieved by a fine control of their secretion. Noteworthy, most of the effectors are synthesized in post-exponential growth phase, similar to the transmissive phase of *L. pneumophila* infectious cycle, and thus available for secretion when the bacteria contact with the host cell [20, 31]. Nevertheless, proteomics showed that few effectors (24/91) are exclusively synthesized during the exponential phase, similar to the replicative phase of the bacteria inside the LCV [31], and consequently would be secreted later than the transmissive phase-effectors. Such control of effector synthesis impacts on the accurate effector delivery, as exemplified by the replicative phase-effector MvcA (Lpg2148) which is detected in the cell 5 h after the transmissive phase-effector MavC, thus allowing the deubiquitinase activity of MvcA to reverse MavC-mediated UBE2N ubiquitination [31, 32]. However, this transcriptional control is not relevant for effectors available upon contact with the host cell and/or synthesized throughout the *Legionella* cycle, thus suggesting that control of effector secretion could also rely on the control of the secretion activity and/or of the substrates recognition by the Icm / Dot system itself.

Effector recognition by the coupling complex DotM protein requires a C-terminal signal of less than 30 amino acids with both short polar amino acids and a glutamate-rich region (E-box) retrieved in some effector sequences [10, 33–35]. Alternatively, some effectors are dependent of the IcmSW chaperone proteins that would scan the Icm/Dot system environment to maximize substrate capture [10] and could recognize other effector motif [36]. Recently, the atomic structure of the fully-assembled *Legionella* T4 coupling complex revealed a DotLMNYZ hetero-pentameric core complex associated with the flexible IcmSW-LvgA chaperone module, thus providing mechanistic models for recruitment and delivery of two types of effectors, the IcmSW-LvgA-dependent class and the E-box one [37, 38]. However, as suggested by the analysis of the secretion determinants of SidJ effector, multiple mechanisms can regulate the export of some effectors [39]. The secretion of SidJ is mediated by dual signal sequences that include a conventional C-terminal domain needed for the secretion at early points of infection, and an internal motif efficient at later time points [39]. Both translocation signals and dependence on effector chaperone might modulate the translocation efficiency of given effectors and ultimately their timing of delivery to the target cell. Yet, the translocation dynamics of effectors, and the related putative control of the Icm / Dot machinery itself have not been thoroughly investigated.

Here, we sought to gain information about the dynamics of the secretion activity of Icm/Dot machinery. To this end, we have developed a kinetic translocation assay, which enables monitoring over several hours of the translocation of Icm/Dot substrates constitutively expressed. We here describe that effectors display specific translocation patterns, independently of their expression control, that are remarkably consistent with their function on specific cellular targets in the host. We analyze the contributing factors and provide evidence for an unexpected role of c-di-GMP signaling in orchestrated effector translocation by the Icm/Dot system.

## Results

### Kinetic assay of effector translocation reveals distinctive and effector-specific profiles

We sought to analyze the temporal activity of the Icm/Dot system in the initial phase of monocyte infection during which the translocation of effectors determines the fate of the bacterium. In order to directly compare the translocation dynamics of multiple effectors, we adapted the *β*-lactamase translocation reporter system [40] that contributed to the identification of a significant part of the 300 Icm/Dot substrates [4]. Effector proteins fused to the C-terminus of the TEM-1 *β*-lactamase and constitutively produced from a Ptac promoter, are detected in host cells by the cleavage of the *β*-lactam ring of CCF4, a fluorescent substrate of TEM-1 that accumulates in eukaryotic cells. In typical end-point translocation assay, CCF4 is added 1h post-infection to quantify the ratio of cleaved/intact *β*-lactamase substrate (emission ratio 460/530 nm). This method provides a quantitative measure of the effector fusion translocated into the host cell, but it is limited to a single time point. A live kinetic assay using CCF4-preloaded cells provided a more dynamic image of effector translocation, but the observable timeframe was limited to less than 90 minutes as a consequence of CCF4 leakage from monocytes [19]. Here, we developed a multiple end-point assay to follow the level of translocated effector over an extended time frame. We took advantage that protonophores, such as carbonyl cyanide m-chlorophenyl hydrasone (CCCP), completely inhibits the Icm/Dot T4SS activity [19], allowing us to freeze translocation at different time points before addition of CCF4. To test this method, a TEM-LepA protein expressing fusion was introduced in *L. pneumophila* Lens and used to infect U937-derived phagocytes. The sensitivity of the assay was assessed by infecting monocytes with an increasing number of bacteria expressing the translocated TEM-LepA fusion protein (Fig. S1, A and B). We have chosen a Multiplicity Of Infection (MOI)=20 as a good compromise between sensitivity and linearity. The concentration of the protonophore CCCP to inhibit the Icm/Dot secretion of the TEM-LepA fusion protein was also set up (Fig. S1C). In addition, we have checked (i) the fusion protein was stably produced by bacteria and thus constantly available for secretion all along the infection (Fig.S1DE), (ii) CCCP at 10 μM was effective in stopping Icm/Dot activity up to 240 min post-infection (Fig. S1F) and (iii) the signal due to the ß-lactamase activity of TEM-LepA fusion protein was stable 2h after the addition of CCCP (Fig. S1G).

Consistent with previous observations, we found that the level of translocated LepA effector steadily increases during the first hour to reach a plateau [19]. However, and unexpectedly, a sudden burst of LepA levels is observed at a later time point (∼100 min.) before returning to lower and steady levels (Fig. 1A). As expected, no translocation was detected when cells were infected by the isogenic mutant strain deleted of the *dotA* gene or when TEM-1 was fused to the housekeeping protein Enoyl-acyl CoA reductase FabI (Fig. 1A). We then used this method to analyze the translocation profiles of the effectors LegK4, SidJ, SidM and LegK2. Despite these effectors are constitutively produced and stable (Fig. S2A), each effector showed distinct, and even opposing, translocation profiles (Fig. 1B). For instance, LegK4 levels are maximal at the earliest time point (10 minutes), confirming that Icm/Dot translocation could occur immediately after the host-cell contact, then rapidly decrease to background levels 1h post-infection. SidJ levels slowly but steadily increase over time while SidM levels sharply increase at 90 minutes to then reach a plateau that likely represents a CCF4 substrate-limiting step. Reminiscent of the burst observed for LepA levels at ∼100 minutes, LegK2 levels also showed a burst at 30 minutes before returning to background levels (Fig. 1B). Noteworthy, the rapid decrease in secreted effector level into the host cell is not due to a decrease in effector synthesis by *L. pneumophila*, as demonstrated by constant detection of TEM-LegK2 in bacteria throughout infection (Fig. S1E). Interestingly, the progressive delivery observed for TEM-SidJ is consistent with the gradually immunodetection during the course of the infection of the endogenous SidJ [41], constantly produced [42]. We compared two other types of secretion profiles, those of SidM and LegK2, with the effector immunodetection on the LCV. Endogenous SidM/DrrA had been detected by immunofluorescence microscopy on the early LCV during the first 3 hours post-infection and with maximal association at 1 hour post-infection [22]. Using a similar microscopy-based immunofluorescence method and *L. pneumophila* expressing a HA-tagged SidM, we measured the level of SidM associated to the LCV relative to mCherry-expressing bacteria (Fig. 1C). We confirmed that HA-SidM is found on the LCVs as early as 30 min post-infection. At 75 minutes, HA-SidM was increasingly associated with LCVs to reach a maximum at 90 minutes post-infection (Fig. 1C). The kinetics of translocated levels of TEM-SidM is highly consistent with the dynamics of localization of HA-SidM on the LCV (Fig. 1C, Fig. S3). Similarly, the secretion profile of TEM-LegK2 and the immunodetection of HA-LegK2 on the LCV both reveal the transient presence of LegK2 in the host cell in the early stages of infection, with a maximum peak shifted by only 15 min between the two methods (Fig. S4, A and B), attesting to the relevance of the profiles obtained from the translocation assays.

**Fig. 1.**
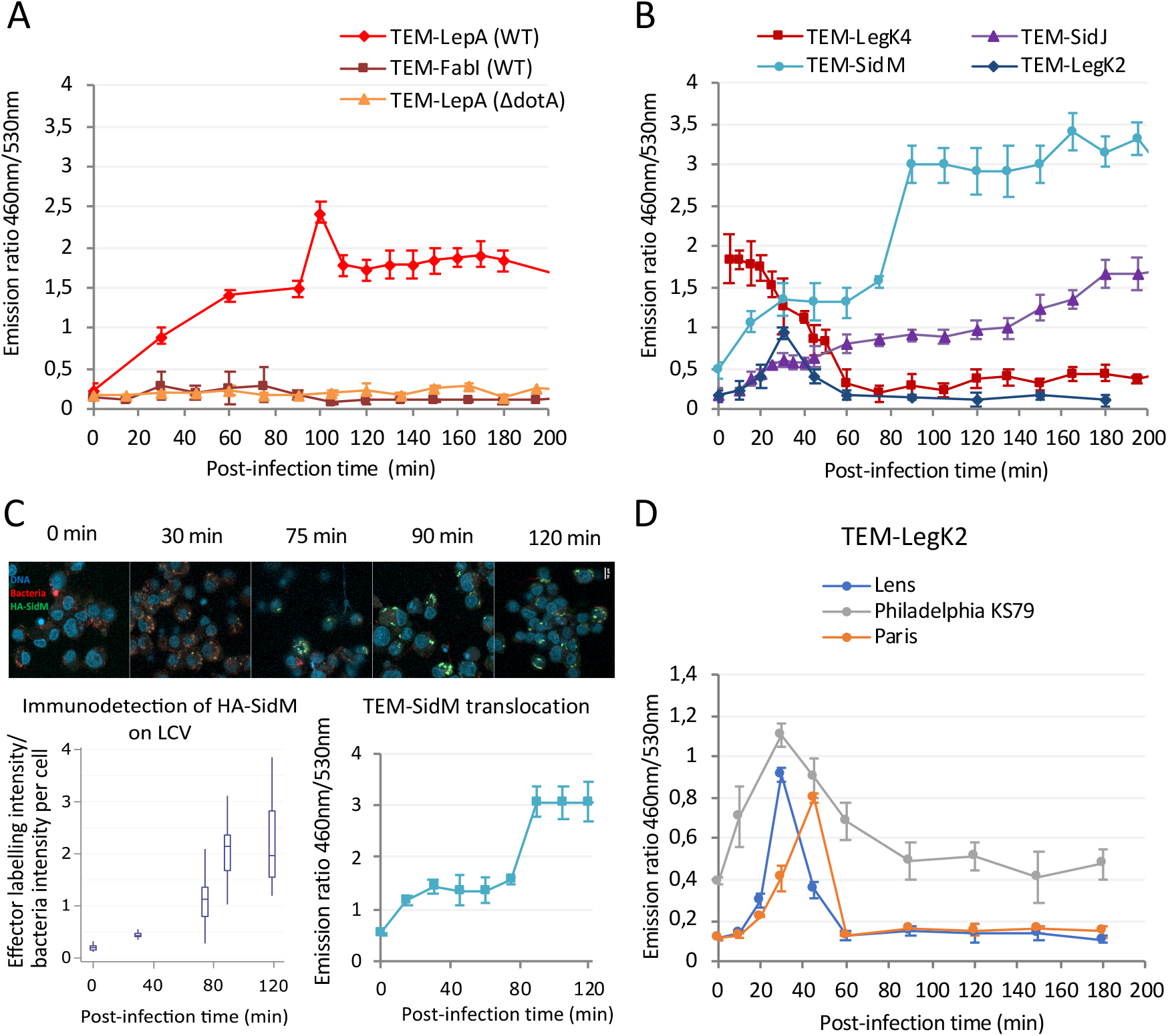
Kinetic assay of effector translocation reveals distinctive and effector-specific profiles. (A) Translocation kinetics of TEM-LepA fusion protein. U937 cells were infected with wild-type (WT) and *ΔdotA* Lens strains harboring a TEM-LepA or a TEM-FabI (non-secreted protein control) expression plasmid (MOI 20). Results are obtained from 3 independent experiments made in triplicates and are presented as means ± SD. (B) Translocation kinetics of TEM-LegK4, TEM-SidJ, TEM-SidM, and TEM-LegK2 in the WT Lens strain. Results are obtained from 3 independent experiments made in triplicates and are presented as means ± SD. (C) Correlation between translocation kinetics of TEM-SidM and the immunodetection of HA-SidM on LCV. *Top panel*, Confocal laser scanning micrographs at 0, 30, 75, 90, and 120 min post-infection of U937 cells infected with HA-SidM and mCherry producing *L. pneumophila* (MOI 50) Nucleus were stained using DAPI (blue), bacteria expressed the fluorescent protein mCherry (red) and HA-SidM were immunolabeled with anti-HA antibodies (green) (Fig. S3 for more details). *Left panel*, quantitative analysis of HA-SidM presence on LCV: the average intensity of HA labeling was reported to the average intensity of bacteria fluorescence for each U937 cell on at less 20 cells per condition. Central box represents the values from the lower to upper quartile (25 th to 75 th percentile) for each condition. The middle line represents the median. *Right panel,* translocation kinetics of TEM-SidM fusion in the WT Lens strain. The results are representative of 3 independent experiments. (D) Translocation kinetic profiles are similar between *L. pneumophila* Lens, Philadelphia KS89 and Paris strains. Translocation kinetics of TEM-LegK2 in *L. pneumophila* Lens, Philadelphia KS89 and Paris strains. These results are representative of 2 independent experiments and are presented as means ± SD.

In addition to profile differences, the maximum of secretion level measured into the host cell is sometimes very different from one effector to another and appears uncorrelated to the level of the effector production (Fig. S2B). This could result either from the intrinsic ß-lactamase activity of the TEM-effector protein fusion in the host cell, or from the intrinsic effector recognition efficiency by the Icm/Dot system. More interestingly, the secretion kinetics are not only consistent with endogenous effector detection but are also almost identical in the Lens, Philadelphia KS79 and Paris strains, which attests to the robustness of the timing of each effector translocation (Fig. 1D and Fig. S5).

Altogether, translocated effector levels can be assessed over the course of several hours and with a temporal resolution that was not previously available. Importantly, the new data draw an unexpected complex picture of translocation profiles characterized by specific timing of increasing levels, which can be either transient or stable.

### Kinetic translocation profiles are consistent with functional consequences

The specific profiles of effector translocation levels suggest that a specific mix of different effectors at a given time could determine the succession of events that follows *L. pneumophila* phagocytosis. One of the best characterized series of events is the biogenesis of the LCV which involves the seemingly sequential action of Icm/Dot effectors targeting the host cell small GTPase Rab1. The GEF activity of SidM/DrrA is known to activate Rab1 on the LCV surface [43, 44], and its AMPylase activity maintains Rab1 in a GTP-linked active form [45, 46]. Even though LidA activity on Rab1 remains more elusive, several structural studies of the LidA-Rab1 complex propose that LidA also contributes to the activation of Rab1 on the LCV, by blocking de-AMPylation by SidD and disrupting the switch function of Rab1, rendering it persistently active on the vacuole [47, 48]. Conversely, the GAP activity of LepB catalyzes the Rab1 GTP-hydrolysis and results in the removal of Rab1 from the LCV surface [22] (Fig. 2A). The kinetics of LepB levels was previously determined but at the low time resolution of 1 timepoint per hour [22, 23]. A time-resolved analysis of the translocated levels of the TEM-SidM, TEM-LidA and TEM-LepB fusions was performed as described above. Similar to SidM, the two other translocated proteins display a pattern with relatively low levels followed by a steady increase to reach a CCF4-limiting plateau (Fig. 2B). However, each effector is characterized by specific timing of increasing levels. As described above, translocated SidM is detected as early as 30 minutes and its levels accumulated quickly, reaching a plateau at 90 min post-infection. LidA follows a similar pattern but significantly delayed. LepB levels rise sharply nearly 1 hour after those of SidM to slowly reach a plateau at 3 hours post-infection (Fig. 2B). Strikingly, the accumulation kinetics of these effectors is in agreement with the function of the corresponding effectors during infection. SidM would be translocated first to activate the Rab1 GTPase on the LCV, followed by LidA that would promote this Rab1 activation. Finally, LepB would be the last translocated effector to remove Rab1 from the LCV, thus terminating Rab1-dependent ER recruitment on the LCV. This experiment supports that the complex orchestration of effector actions relies, at least in part, on the defined timing of translocated levels of some of the numerous Icm/Dot effectors into the host cells.

**Fig. 2.**
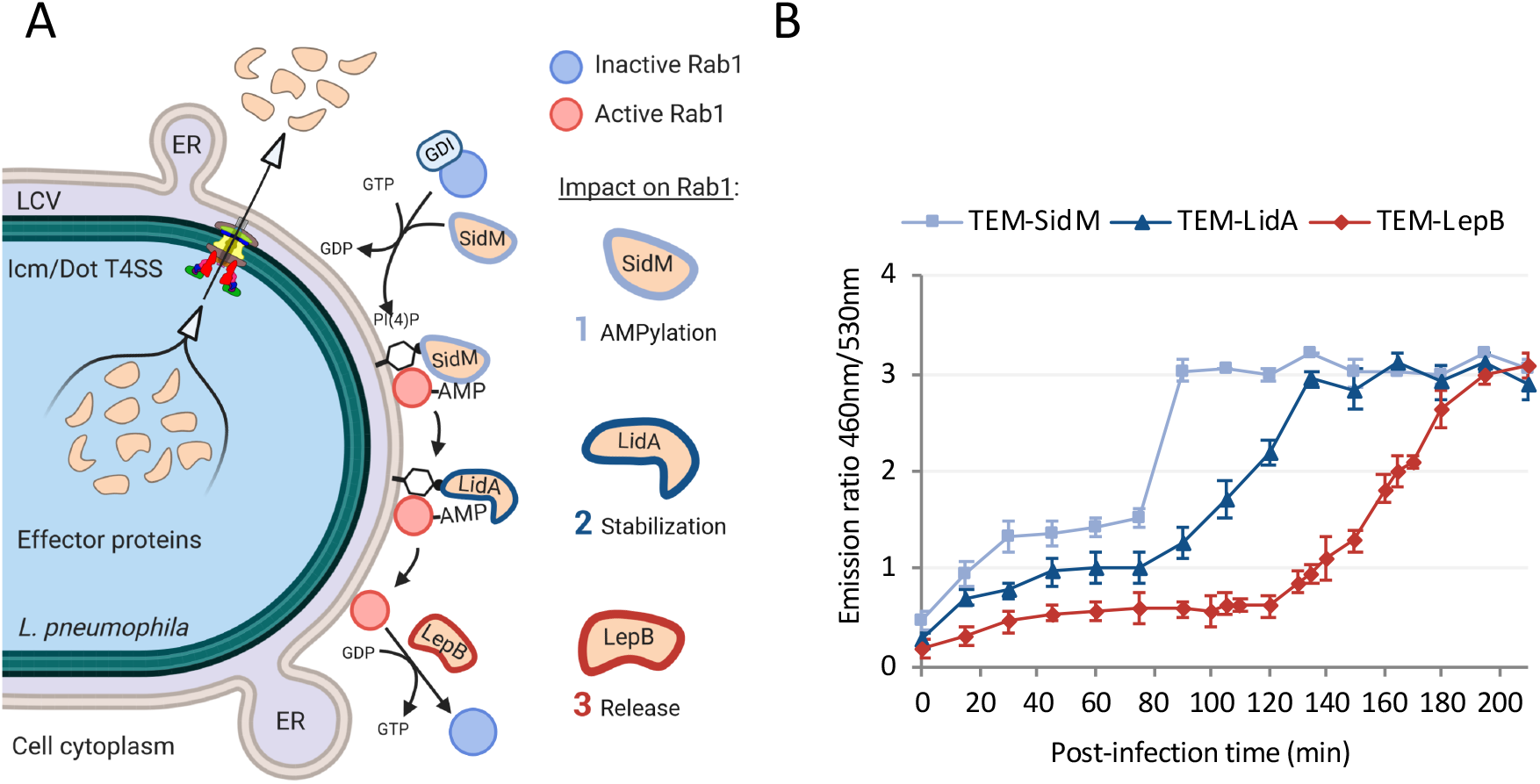
Kinetic translocation profiles are consistent with functional consequences. (A) Sequential action of SidM, LidA and LepB controls the Rab1 small GTPase activation during LCV biogenesis. Lipids remodeling of the LCV membrane ([PI(4)P] enrichment) promote the binding of some *Legionella* effectors on the LCV. Several of them control the recruitment and activation of the Rab1 small GTPase at the surface of the LCV, notably SidM (GEF and AMPylase), LidA, and LepB (GAP). GDI: dissociation inhibitor; ER: Endoplasmic reticulum; LCV *Legionella* containing vacuole. (B) Translocation kinetics of TEM-SidM, TEM-LidA, and TEM-LepB in the WT Lens strain. Results are obtained from 3 independent experiments made in triplicates and are presented as means ± SD.

### Robust timing of effector translocation, independently of effector concentration

The finding that various effectors begin to accumulate in the host cell with a specific timing suggests that the translocation apparatus exerts a control on the translocation process. Such control has been previously documented for type III secretion systems (T3SS). The main factors contributing to the timing of effector translocation are the relative concentration of effectors in the bacterial cytoplasm and the involvement of chaperones [49]. The timing at which various *L. pneumophila* effectors begin to accumulate in the cell is strikingly different despite the fact that they are all ectopically, constitutively and stably expressed (Fig. S2A). Yet, we could not rule out that small difference in effector concentration dictates the timing of translocation. To directly test this hypothesis, we monitored the translocation profiles of TEM-LepA and TEM-LegK2, expressed at increasing levels from an IPTG-inducible promoter. From 0.5 µM to 500 µM IPTG, increased expression levels of TEM-LepA or TEM-LegK2 result in increasing levels of translocated effector (Fig. 3A and B). However, the secretion profile of each effector is conserved, with specific bursts and maximal levels conserved (Fig. 3A and B). Consistently, the secretion kinetic profiles of TEM-LepA and TEM-LegK2 are also conserved independently of the MOI used during the assays (Fig. 3C and D). Thus, the kinetic profiles of LepA and LegK2 translocation are largely unaffected by their expression levels, disproving the hypothesis that the timing of effector translocation is controlled by effector expression levels.

**Fig. 3.**
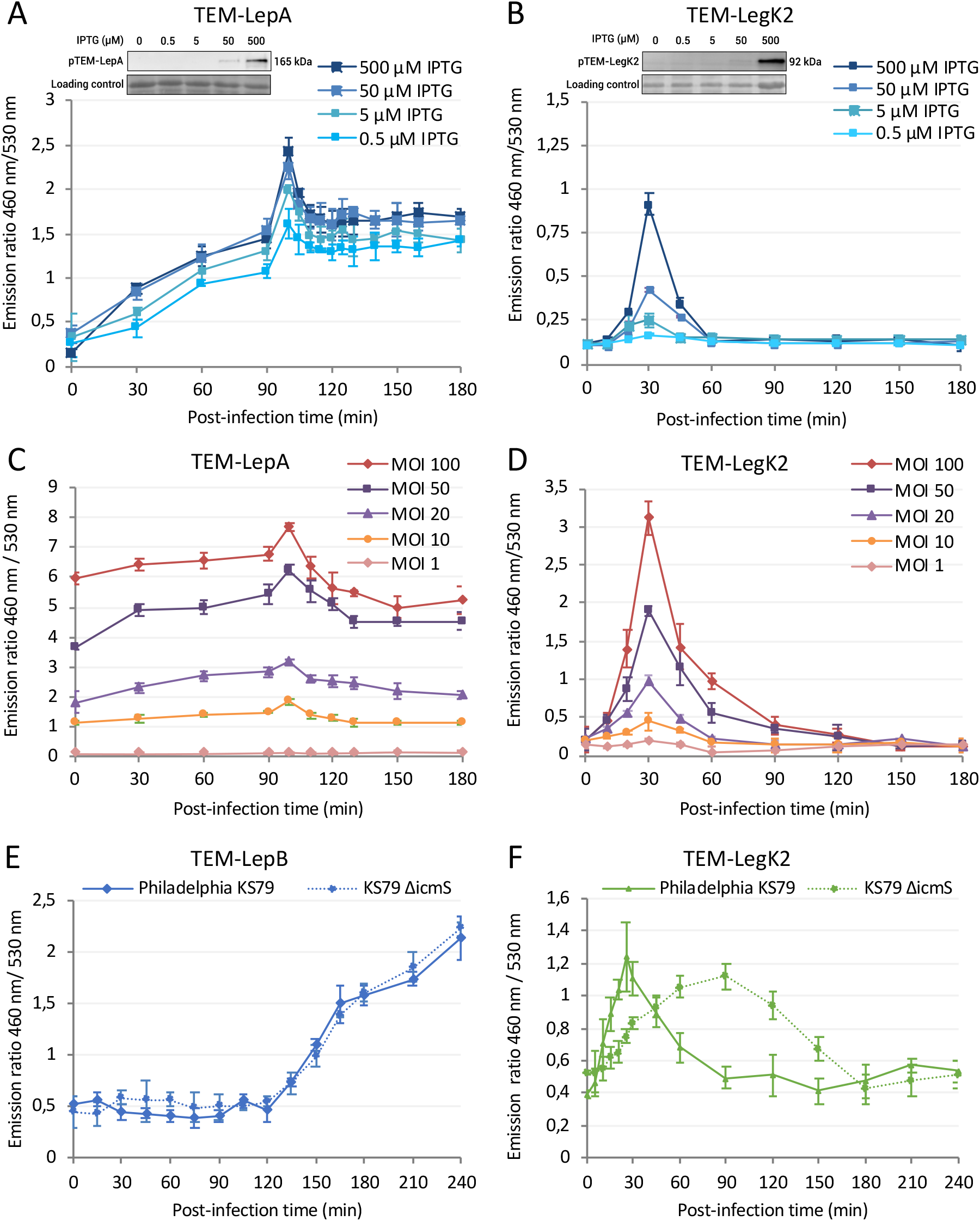
Robust timing of effector translocation, independently of effector concentration and chaperones activity. (A and B) Translocation kinetics of TEM-LepA (A) and TEM-LegK2 (B) fusion proteins during infection of U937 (MOI 20), after induction of their expression by a range of IPTG concentrations corresponding to different levels of synthesis assessed by Western blot with anti-TEM (top panel). (C and D) Translocation kinetics of TEM-LepA (C) and TEM-LegK2 (D) fusion proteins during infection of U937 cells at various MOI. (E and F) Translocation kinetics of the IcmS-independent TEM-LepB (E) and the IcmS-dependent TEM-LegK2 (F) fusions in KS79 *L. pneumophila* Philadelphia and in the *ΔicmS* mutant strain. All the results are representative of 2 independent experiments and are presented as means ± SD.

Translocation of some effectors has been described to be dependent on the chaperone IcmS-IcmW while translocation of other effectors, does not require IcmS-IcmW [13]. Thus, recognition by the IcmS-IcmW may determine the timing and efficiency of effector translocation. We tested this hypothesis by monitoring the secretion of TEM-LepB, known to be translocated independently of IcmS-IcmW [50], and TEM-LegK2, whose we revealed a chaperone-dependent translocation, by the strain derived from Philadelphia KS79 and its isogenic mutant Δ*icmS*. Expectedly, the absence of IcmS had no impact on the kinetics of LepB translocation (Fig. 3E). By contrast, the absence of IcmS had a major impact on LegK2 levels which rose less rapidly, remained lower but lasted for an extended time (Fig. 3F). Rather than delivering a short burst of LegK2, the Δ*icmS* seemed to translocate the same amount of LegK2 but over a longer period of time, consistently with a chaperon-dependent translocation (Fig. 3F). However, and importantly, the lack of IcmS impacted translocation efficiency of LegK2 but it had no impact on the precise time at which translocated levels of LegK2 begin to rise (at ∼5 minutes). Thus, the Icm/Dot system starts translocation of effectors at a defined timing that does not depend on effector concentration or chaperone activity. The mechanism that controls the timing of translocation of each effector is unknown but may be related to the properties of the effector protein (E-box, folding rate, stability) and how each effector interacts with the Icm/Dot system (affinity). Nevertheless, timing of effector translocation may also result from intrinsic and/or external modulations of the activity of the Icm/Dot system.

### Timing of Icm/Dot effector translocation depends on c-di-GMP signaling

We previously reported that three c-di-GMP-metabolizing enzymes directly contributed to the ability of *L. pneumophila* Lens strain to infect both protozoan and mammalian cells. Mutants with deletions of the corresponding genes (*lpl0780*, *lpl0922* and *lpl1118*) are not defective for planktonic multiplication or entry into the host cell, but were partially defective for Icm/Dot-dependent processes such as escape of the LCV from the host degradative endocytic pathway and efficient endoplasmic reticulum recruitment to the LCV, and consequently are significantly delayed for intracellular replication [51]. A snapshot of effector translocation, 1 hour post-infection, revealed effector-specific alterations in translocation efficiencies. While some effectors appeared unaffected, others appeared translocated less efficiently while, even more surprisingly, others appeared over-translocated [51].

Interestingly, while no significant phenotype of intracellular replication had been revealed in the Philadelphia strain [52], we here observe that the intracellular replication of the *lpp0809* deletion mutant were importantly delayed in the Paris strain, even more than in the *lpl0780* mutant in Lens strain, suggesting a conserved function in the Lens and Paris genetic backgrounds (Fig. 4A). The sequences of Lpl0780 and Lpp0809 are strongly conserved (97 % identity, 99 % similarity) and as expected, the *lpl0780* gene can rescue the phenotype of the Δ*lpp0809* strain (Fig. 4B). Then, we sought to address the potential role of the diguanylate cyclase (DGC) Lpl0780/Lpp0809 in the control of effector translocation, reasoning that difference in effector translocation efficiencies in the previous snapshot actually resulted from alterations in the timing of effector translocation. Hence, we analyzed the translocation kinetics of the TEM-LepA effector in the Lens and Paris strains deleted of the gene *lpl0780* or *lpp0809*, respectively (Fig.4C and D). Interestingly, the burst of LepA levels appeared delayed by nearly 1 hour in the both *lpl0780* and *lpp0809* mutants. Similarly, translocation of the LegK2 effector is delayed in the *lpl0780/lpp0809* mutants. More precisely, despite the overall translocation profiles remained nearly identical, initiation of translocation occurs with a 40 minutes delay (Fig. 4E and F). While the introduction of the *lpl0780* gene on a plasmid restored the normal delivery of the LepA and LegK2 effectors in the Δ*lpl0780* strain, a mutation in the catalytic GGDEF domain of the Lpl0780-encoding gene [51] abolished the rescue of the Δ*lpl0780* strain (Fig. 4C and E). Altogether, the data support a role of the c-di-GMP synthesizing activity of the DGC Lpl0780/Lpp0809, and subsequently of c-di-GMP signaling, in the timing of effector translocation. Noteworthy, the overexpression of *lpl0780* in the wild-type (WT) Lens strain has no significant impact on the both profiles (Fig. S6AB), while the overproduction of the phosphodiesterases Lpl0220 and Lp1118, known to cause a significant decrease in the concentration of c-diGMP in *Legionella* [52], considerably reduces the secretion of TEM-LegK2 and also of TEM-LepA in the case of the overproduction of Lpl0220 (Fig. S6CD). These results suggest that a basal and accurate endogenous level of c-di-GMP is essential to Icm/Dot secretion.

**Fig. 4.**
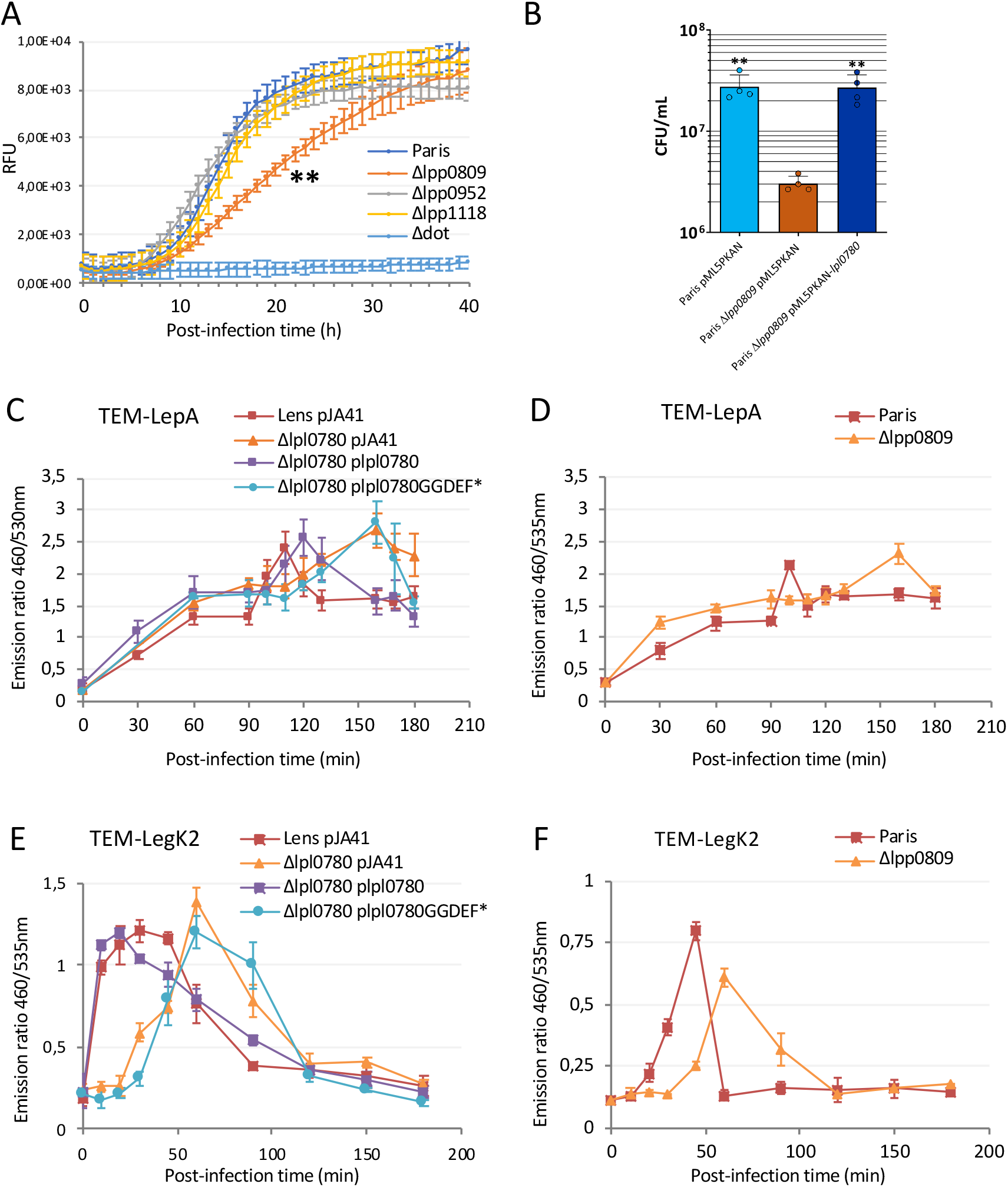
Timing of Icm/Dot effector translocation depends on c-di-GMP signaling. (A) Phenotype conservation between Δ*lpp0809* (Paris derivative) and Δ*lpl0780* (Lens derivative) strains. Intracellular replication of mCherry-producing Paris derivative strains (WT, Δ*lpp0809,* Δ*lpp0952*, Δ*lpp1114* and Δ*dotA* strains) in *A. castellanii* (MOI 10) were monitored by fluorescence and compared to the previously published phenotypes of Lens derivative strains (Δ*lpl0780*, Δ*lpl0922* and Δ*lpl1118*) [51]. Significance was calculated in comparison to the Δ*lpp0809* strain with non-parametric Mann-Whitney test (** => p ≤ 0,002). The results are representative of 3 independent experiments. (B) The *lpl0780* gene rescues the phenotype of the Δ*lpp0809* Paris strain. CFU from U937 cells infected by Paris strains expressing *lpl0780* gene for 48 hours (MOI 10). Significance was calculated in comparison to the Δ*lpp0809* strain with non-parametric one-way ANOVA (** => p ≤ 0,002). The results are representative of 2 independent experiments. (C and D) Translocation kinetics of TEM-LepA fusion protein in WT, Δ*lpl0780* and complemented Lens derivative strains (C) or in WT, Δ*lpp0809* Paris derivative strains (D). These results are representative of 2 independent experiments and are presented as means ± SD. (E and F) Translocation kinetics of TEM-LegK2 fusion protein in WT, Δ*lpl0780* and complemented Lens derivative strains (E) or in WT, Δ*lpp0809* Paris derivative strains (F). These results are representative of 2 independent experiments and are presented as means ± SD.

### The DGC Lpl0780/Lpp0809 most likely acts at a post-translational level at the bacterial poles

To go further on the Icm/Dot secretion control by c-di-GMP, we addressed the question whether the DGC Lpl0780/Lpp0809 could impact the expression of genes encoding components of the secretory apparatus and /or effectors. RNA sequencing showed that only 12 genes were significantly differentially expressed (*P <* 0.01 and log_2_ fold change of > 1 or < −1) between *Δlpl0780* strain and the WT Lens strain (table S4). Noteworthy, the fold changes remain quite weak and none of these genes can be connected to T4SS machinery or effector expression, thus suggesting a post-transcriptional control. Among the described modes of action of c-di-GMP [53], we then favored the hypothesis of a post-translational control by synthesis of c-di-GMP close to the Icm/Dot secretion machinery by the DGC Lpl0780/lpp0809. Given that the secretion apparatus was recently described to be located at the bacterial poles [17], we tested this hypothesis by determining the cellular localization of Lpp0809-sfGFP fusion protein expressed from a plasmid or from the chromosome (Fig. 5 and Fig. S7). Importantly, in both cases the Lpp0809-sfGFP fusion proteins are stable and functional, as demonstrated by their ability to restore intracellular replication of the *Δlpp0809* strain (Fig. 5A and B). The plasmid-encoded version of Lpp0809-sfGFP localizes at both poles in 30% of the stationary phase cells, and sometimes in the form of foci along the bacterium (Fig. 5C-E). Moreover, internal foci are observed for larger cells (Fig. 5D), as described for the machinery component DotF proposed to be targeted to the pole at the midcell [17]. To check that the Lpp0809 polar localization is not due to overexpression of the fusion protein or to the sfGFP tag, localization of Lpp0809-sfGFP and 4xHA-Lpp0809 fusion proteins synthesized from chromosome was established. In half of the bacteria, the chromosome-encoded version of Lpp0809-sfGFP is also detectable, despite weak fluorescence, at the poles (Fig. S7) with a polarity score similar to that of T4SS components outside the complex core, such as DotB or DotL [15]. Bipolar localization is also observed by immunodetection of 4xHA-Lpp0809 (Fig. S7D) and the fusion protein is observed close to the coupling complex protein DotN in about 50% of bacteria (Fig. 5F). Interestingly, polar localizations of Lpp0809 and DotN are independent of each other (Fig. S8), which may suggest that polar localization of the DGC is not dependent on a direct interaction with this secretion machinery component. Together, these data suggest that the DGC Lpl0780/Lpp0809 could modulate the local pool of c-di-GMP near the Icm/Dot machinery, thus controlling directly or indirectly the initiation of effector translocation.

**Fig. 5.**
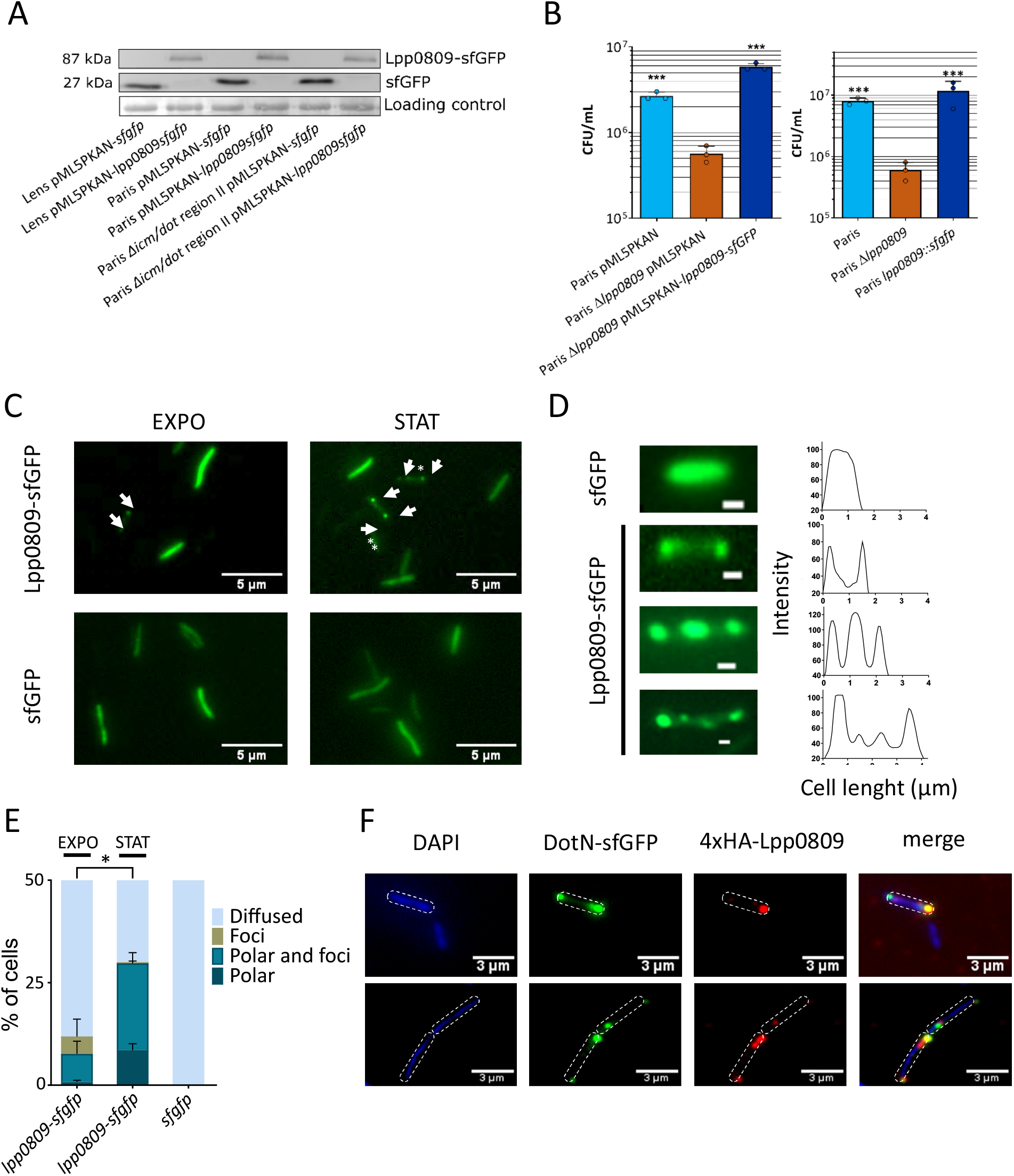
The DGC Lpl0780/Lpp0809 most likely acts at the bacterial poles. (A) Lpp0809-sfGFP fusion protein is stable. Anti-GFP immunoblots analysis of protein extracted from pML5PKAN derivative containing Lens strains expressing sfGFP or Lpp0809-sfGFP. The results are representative of 2 independent experiments. (B) Lpp0809-sfGFP fusion protein is functional. Intracellular replication of Paris strains expressing the pML5PKAN-*lpp0809-sfgfp* fusion (left panel) or a chromosomal copy of *lpp0809-sfgfp* fusion inserted at the *lpp0809* locus (right panel) in U937 cells was measured after 48 hours (MOI 10). These results are representative of 3 independent experiments and are presented as means ± SD. Significance was calculated in comparison to the Δ*lpp0809* depleted strain with non-parametric one-way ANOVA (*** => p ≤ 0,001). (C) Localization of recombinant Lpp0809-sfGFP in Paris strain expressing the pML5PKAN-*lpp0809-sfgfp* fusion, collected in exponential and stationary phases. Cells with polar localization (arrowheads) and foci localization (asterisk) of Lpp0809-sfGFP are indicated (scale bar = 5µm). (D) ImageJ analysis of fluorescence intensity along the axis of representative cells (scale bar = 0,5µm). (E) Percentage of cells displaying cytoplasmic, foci or polar localization of Lpp0809-sfGFP (n ≥ 200 bacteria in exponential and stationary phases). The results are representative of 2 independent experiments. (F) Localization in Paris strain of recombinant DotN-sfGFP (green), 4xHA-taggedLpp0809 (red) and DAPI staining (blue) (scale bar = 3µm) (n ≥ 200 bacteria).

## Discussion

We developed a kinetic translocation assay that allows a fine monitoring of translocation of Icm/Dot substrates constitutively produced by *L. pneumophila*, and consequently of the secretion activity of the T4SS, independently of the expression level of the effectors.

It is based on the *β*-lactamase (TEM) translocation reporter system combined with the effect of the protonophore CCCP which, added at various time points, collapses the proton gradient and thus stops the translocation process [19]. Its temporal resolution is dictated by the timing of addition of CCCP which may be modulated depending on the time window investigated. The CCCP-combined *β*-lactamase assay has a lower resolution power than the previously published real-time *β*-lactamase assay but allows monitoring on larger time frame. Although monitoring could be extended to several hours, a 3-hours window was sufficient to observe major difference in kinetic translocation profiles of effectors [49]. The physiological relevance of the translocation kinetics obtained with the CCCP-combined *β*-lactamase assay was validated by demonstrating that the time-dependent SidM and LegK2 translocation into the host cell was consistent with the immunodetection of these effectors on the LCV during infection. Thus, we observed that SidM accumulated into the host cells from 75 min to 200 min, consistently with the western-blot detection of SidM in the host cell between 1h to 3h post-infection [23]. Likewise, our assay confirms that when SidJ is constitutively expressed, the amount of translocated SidJ increases steadily after phagocytosis, as observed previously for the endogenous SidJ effector whose protein level is constant throughout the bacterial growth cycle [42]. Importantly, this consistency emphasizes that temporal control of the presence of an effector in the host cell is primarily controlled by the regulation of the effector docking / translocation by the secretion machinery, rather than by the control of effector synthesis. Nevertheless, in some cases, such as MavC and MvcA which we have both observed to be translocated at 25 min post-infection when constitutively expressed (Fig S2B) while the ubiquitin deamidase of MvcA is known to reverse the noncanonical ubiquitination mediated by MavC, the temporal regulation is mainly achieved by differential expression of the genes at different stages of the intracellular life cycle of *L. pneumophila* [20].

We observed rapid and early translocation of some effectors (e.g LegK4) consistent with the fact that the Icm/Dot secretion apparatus is already assembled before the contact with the host cell [14, 17]. However, late translocation of some effectors (e.g LepB) strongly supports the occurrence of other signaling mechanisms to control their translocation. Significantly, we demonstrated that the early translocation is not due to the effector recognition by the chaperone protein IcmS. However, the IcmS protein is essential for the translocation rate, consistently with the model proposed by Meir and *al.* [10]. Importantly, we also showed that the timed delivery of an effector is not dependent on its concentration and synthesis. Rather, the timing of effector translocation seems to be determined by the intrinsic properties of each effector protein (E-box, folding rate, stability). As exemplified by the Icm/Dot effectors which target the host cell small GTPase Rab1, these properties may dictate a translocation timing that determine the effectors sequential actions in the target cell.

Besides, we highlighted that in addition to the control of translocation initiation, the decrease in levels of translocated effector is also regulated. Translocated levels of some effectors quickly decrease after the beginning of the translocation (e.g LegK2), while that of others is maintained for a long time period, until 4h post-infection (e.g SidJ). Consistently, it has emerged that *L. pneumophila* is able to achieve temporal regulation of an effector using the ubiquitin-proteasome system [54]. Indeed, after establishing its replicative niche, the *L. pneumophila* effector SidH is degraded by the host proteasome. Most remarkably another effector protein LubX is able to mimic the function of a eukaryotic E3 ubiquitin ligase and polyubiquitinates SidH, targeting it for degradation [29, 55]. Together, these data clearly demonstrate that *L. pneumophila* temporally controls the function of Icm/Dot effectors inside host cells by (1) controlling effectors synthesis, (2) fine-tuning the initiation of secretion, (3) controlling the rate of effector translocation, and (4) modulating the half-life of some effectors after their delivery into the cells. Additionally, it is clear that the function of these effectors is also controlled spatially by their addressing to the appropriate host cell compartment [56, 57].

We here provide evidence that Icm/Dot secretion is controlled by c-di-GMP signaling, as inactivation of some c-di-GMP synthetizing DGC, or overproduction of some c-di-GMP degradation enzymes, results in significant changes in the rate and timing of some effector translocation. In particular, we demonstrated that a c-di-GMP-synthesizing enzyme, namely the DGC Lpl0780/Lpp0809, significantly contributes to the triggering of effector translocation. We propose that this enzyme most likely exerts a post-translational control of the T4SS machinery rather than a control of its synthesis. Given its localization at the bacterial pole with the T4SS machinery and by comparison with previously described post-translational controls of other bacterial secretion systems (except the T4SS) by binding of c-di-GMP [58–63], it is tempting to hypothesize that a “high specificity local c-di-GMP signaling” [64] could modify the interactions between the coupling protein complex, the chaperones and others unknown partners to finely orchestrate loading of the 300 effectors on the Icm/Dot in coherence with their role during the infection. Even though our results do not yet establish the detailed mechanism involved, they reveal an unexpected c-di-GMP dependent control of the type IV secretion that could complete the recent models of capturing and delivery of the huge number effectors of *Legionella* [10].

## Materials and Methods

### Bacterial strains, culture, and growth conditions

Bacterial strains and cell line used in this study are summarized in table S1. *Legionella pneumophila* strains were grown at 30°C or 37°C either on buffered charcoal yeast extract (BCYE) agar, *Legionella* growth liquid medium (LGM)[51] or ACES (*N*-(2-acetamido)-2-aminoethanesulfonic acid) Yeast Extract medium (AYE). Each media was supplemented with chloramphenicol 5 µg.ml^-1^ kanamycin 50 µg.mL^-1^ gentamicin 5 µg.ml^-1^ and Isopropyl-β-d-thiogalactopyranoside (IPTG) 1 mM or 500 µM when appropriate. *Escherichia coli* strains were grown at 37°C in LB liquid medium or LB-agar medium supplemented with chloramphenicol 5 µg.ml^-1^ or kanamycin 50 µg.mL^-1^ when appropriate. U937 monocyte cells were cultivated at 37°C in 5 % CO_2_ in RPMI 1640 medium (ThermoFisher Scientific) supplemented with 10 % heat-inactivated foetal calf serum (HyClone^TM^). The differentiation of U937 monocytes into macrophages is conducted during 2 days at a phorbol 12-myristate 13-acetate (PMA) concentration of 100 ng.ml^-1^.

### General molecular biology techniques

The plasmids and primers used in this study are shown in tables S2 and S3, respectively. Standard techniques were used for nucleic acid cloning and restriction analysis. SLIC cloning was performed following the procedure described by Li & Elledge [65]. Restriction enzymes, T4 DNA ligase and T4 DNA polymerase were purchased from New England Biolabs. Plasmid DNA from *E. coli* was prepared by rapid alkaline lysis [66]. PCR amplifications were carried out with Phusion polymerase (Finnzymes) or PrimestarMAX (Takara Clontech) as recommended by the manufacturers. Purification of DNA fragments from agarose gels for subcloning was carried out with a QIAquick gel purification kit (Qiagen). Competent cells of *E. coli*, obtained after chemical treatment with calcium chlorure (CaCl_2_), are transformed by thermal shock with 1 ng of plasmid DNA. *L. pneumophila* is transformed by electroporation (2.4kV, 100Ω and 25µF) with 1 ng of plasmidic DNA.

### Gene inactivation in *L. pneumophila*

Scar-free mutants of the *lpp0809* gene and the *icm/dot* region II were constructed in two steps, taking advantage of a homologous-recombination strategy with the counter-selectable *mazF-kan* (MK) cassette [67].The 2000 bp upstream and downstream regions of the gene of interest were amplified by PCR with primers carrying 30-nt sequences (upstream region: P1-P2 primer pair; downstream region P3-P4 primer pair) complementary to the MK cassette. The upstream and downstream regions were assembled to the MK cassette (amplified from plasmid pGEM-*mazF-kan* with MazFk7-F/MazF-R primers) by PCR overlap extension and used for natural transformation. Once transformed, the transformants are selected on CYE + kanamycin and counter-selected for sensitivity to IPTG; Integration of the cassette at the correct locus was verified by PCR. To obtain scar-free mutants, a second step was performed as follows. Upstream and downstream regions of *lpp0809* have been amplified with primers carrying a 20-nt tail sequences corresponding to the 3’ end of upstream or downstream region (upstream region: P1-P5 primer pair; downstream region P4-P6 primer pair), respectively. These PCRs were assembled by PCR overlap extension and used to transform the previous *Δlpp0809::mazF-kan* strain. Transformants were selected on CYE + IPTG and tested for sensitivity to kanamycin. Scar-free deletion of the *lpp0809* gene and the *icm/dot* region II was verified by PCR and sequencing.

*icmS*^-^ derivative of KS79 strain was obtained by introducing the *icmS3001*::*Kan* mutation of GS3001 strain [68] by natural transformation.

### Chromosomic *lpp0809-sfgfp* gene fusion

Chromosomic *lpp0809-sfgfp* gene fusion were constructed in two steps. The 2000 bp upstream and downstream regions of the *lpp0809* gene terminal codon were amplified by PCR with primers carrying 30-nt sequences (upstream region: P1-P2 primer pair; downstream region P3-P4 primer pair) complementary to the MK cassette. The upstream and downstream regions were assembled to the MK cassette (amplified from plasmid pGEM-*mazF-kan* with MazFk7-F/MazF-R primers) by PCR overlap extension and used for natural transformation. Once transformed, the transformants are selected on CYE + kanamycin and counter-selected for sensitivity to IPTG; Integration of the cassette at the correct locus was verified by PCR. To obtain *lpp0809-sfgfp* mutants, a second step was performed as follows. Upstream and downstream regions of *lpp0809* have been amplified with primers carrying a 20-nt sequences corresponding to either a DNA linker encoding Arg-Tr-Gly-Gly-Ala-Ala (5’-AGAACTGGTGGTGCTGCT-3’) or the 3’ end of *sfgfp* gene (upstream region: P1-P5 primer pair; downstream region P4-P6 primer pair), respectively. These PCRs were assembled by PCR overlap extension and used to transform the previous *Δlpp0809*::*mazF-kan* strain. Transformants were selected on CYE + IPTG and tested for sensitivity to kanamycin. Scar-free deletion of *lpp0809* was verified by PCR and sequencing.

### Analysis of the TEM-effector and sfGFP protein fusion by Western-blot

*L. pneumophila* strains expressing TEM-effector fusions (from pXDC61 derivative plasmids) were grown in 2 mL of AYE medium containing chloramphenicol and 500 µM IPTG at 37°C overnight. *L. pneumophila* strains expressing sfGFP protein fusions (from pML5PKAN derivative plasmids) were grown in 2 mL of AYE medium chloramphenicol and at 37°C overnight. Stationary-phase bacteria were washed 3 times with 1 mL of distilled water and then disrupted with 0.25 mm diameter glass beads in a FastPrep-24 instrument (MP Biomedicals), using six 30-seconds pulses at a speed of 6.5 m/s at room temperature, separated by 5 min intervals on ice. Cell debris were pelleted by centrifugation (13000 x *g*, 20 min). The protein concentrations of intracellular extracts were determined, and equivalent protein quantities were loaded and separated by SDS-PAGE. Loading control was revealed by red Ponceau treatment of the membrane. Then, proteins were transferred on Optitran cellulose nitrate membrane BA-S 85 (pore size 0.45 µm, Schleicher & Schuell) and membranes were saturated with TBS (Tris 100 mM, NaCl 150 mM, pH8) supplemented with non-fat dry milk 5%, incubated with a 1/10000 diluted anti-TEM rabbit serum (Faucher *et al.* 2011) or 1/5000 anti-GFP serum. Detection was performed with anti-rabbit IgG-peroxydase antibodies (A0545, Sigma-Aldrich). Immunoblots are revealed with Pierce SuperSignal West Pico Chemiluminescent substrates (ThermoScientific).

### TEM translocation assays

U937 cells grown in RPMI containing 10% fetal calf serum (FCS) were seeded in black clear-bottom 96-wells plate at 10^5^ cells per well in presence of 100 ng/ml of Phorbol-12-myristate-13-acetate (PMA) to differentiate U937 cells in macrophages. *L. pneumophila* strains carrying the various TEM fusions were grown on BCYE plates containing chloramphenicol (for pXDC61 derivatives), and also gentamicin in case of complementation/overproduction assays with pJA41 derivatives. Bacteria were then grown in AYE containing chloramphenicol, gentamicin when appropriate, and 500 µM IPTG for 72 h to induce production of the hybrid proteins. 10 µL of stationary-phase bacteria re-suspended into RPMI at 2.10^8^ cells.mL^-1^ were used to infect U937 cells (MOI = 20). After centrifugation (880 x *g*, 10 min) to initiate bacteria-cell contact, the plate was incubated at 37°C with CO_2_ exchange. At different time points, 10 µM of CCCP were added to block the secretion through the Dot/Icm T4SS. Cell monolayers were then loaded with the fluorescent substrate by adding 20 µl of 6X CCF4/AM solution (LiveBLAzer-FRET B/G Loading Kit, Invitrogen) containing 15 mM Probenecid (Sigma). The cells were incubated for an additional 90 min at room temperature. Fluorescence was quantified on an Infinite M200 microplate reader (Tecan) with excitation at 405 nm (10 nm band-pass), and emission was detected via 460 nm (40 nm band-pass, blue fluorescence) and 530 nm (30 nm band-pass, green fluorescence) filters [69].

### SidM protein localization during U937 infection

U937 cells grown in RPMI containing 10 % fetal calf serum (FCS) were seeded in 24 wells plate onto sterile glass coverslips at 2.10^5^ cells per well in presence of 100 ng/ml of Phorbol-12-myristate-13-acetate (PMA) to differentiate U937 cells in macrophages. *L. pneumophila* strains carrying pHA-SidM-mCherry vector were grown on BCYE plates containing chloramphenicol. Bacteria were then grown in LGM containing chloramphenicol for 21 h at 37°C to reach the stationary phase. 2 hours before use, mCherry and HA-tagged SidM proteins production were induced by addition of 2 mM IPTG. Monolayers were infected at a MOI of 50. The plates were spun at 880 x *g* for 10 min and cells were immediately fixed or incubated for 30, 75, 90 and 120 min at 37°C. Monolayers were fixed with 4% paraformaldehyde and the aldehyde sites were saturated with glycin. The cells were permeabilized with 100% ice-cold methanol then a saturation step was realized with 1% BSA. The coverslips were stained with anti-HA antibodies from Cell Signaling (ID: 3724S) and visualized with Alexa Fluor® 488-conjugated secondary antibody (Molecular Probes, A11034). The cells were visualized with a DAPI labelling. Microscopy was carried out on a laser-scanning confocal microscope (LSM710 Meta, Zeiss).

### Immunodetection of TEM-effector during *Legionella* infection

U937 cells grown in RPMI containing 10% fetal calf serum (FCS) were seeded in 6 well plate at 10^6^ cells per well in presence of 100 ng/ml of (PMA) to differentiate U937 cells in macrophages. Differentiated U937 cells were infected with a TEM-LepA or TEM-LegK2 expressing bacterial strain (MOI= 20). Infected cells were incubated at 37°C and collected at different time point (0, 30, 90, 120 or 180 min) by scraping the cells in the well. The cells were lysed in 5X Laemmli buffer and the lysates were subjected to SDS-PAGE followed by immunoblotting as previously described using an 1/10000 diluted anti-TEM rabbit serum and an 1/10000 diluted anti-actin antibodies (Sigma-Aldricht).

### Immunofluorescence and fluorescence Microscopy of *L. pneumophila*

HA-Lpp0809 proteins were immunodetected by the procedure previously described for *E. coli* [70] and adapted to *L. pneumophila* [17]. Briefly, *L. pneumophila* strains were grown in AYE until the OD600 reached 1-1.5 (exponential phase) or 3.5-5 (stationary phase). 7 µL of a culture of were fixed for 5 min in 200 µL of 80% methanol, washed in PBS, and allowed to adhere to poly-L-lysine–coated microscope slides. A lysozyme solution [3 mg lysozyme/mL in 25 mM Tris·HCl (pH 8.0), 50 mM glucose, 10 mM EDTA] used to permeabilize cells. The permeabilized cells then were incubated with an anti-HA AlexaFluor®488 conjugated primary antibody (1:800) from Cell Signaling (ID: 2350S). The cells were also stained with DAPI to detect DNA and then were viewed using an Evos FL Cell Imaging System from Thermofisher (100 x objective). Recombinant sfGFP labelled proteins were localized by following the same fixation procedure. Slides were viewed using an Evos FL Cell Imaging System from Thermofisher (100x objective). All measurements of fluorescence intensity and cell length were done using ImageJ (National Institutes of Health).

### Accession number

RNA-seq data have been deposited in European Nucleotide Archive database at EMBL-EBI (https://www.ebi.ac.uk/ena) under accession number PRJEB33700.

## Acknowledgments

The authors of this paper would like to acknowledge Christophe Ginevra for his assistance with RNAseq data analysis. This work was funded by the Centre National de la Recherche Scientifique (UMR 5308), the Institut National de la Recherche Médicale (U1111), the Université Lyon 1. **Funding:** This work was performed within the framework of the LABEX ECOFECT (ANR-11-LABX-0042) of the Université de Lyon, within the program Investissements d’avenir (ANR-11-IDEX-0007) operated by the French National Research Agency (ANR). The Ph.D. grants to J.A., C.J., C.M. were provided by the Ministère de l’enseignement supérieur, de la Recherche et de l’Innovation. **Author contributions:** P.D., A.V. and X.C. designed research; J.A., C.J, C.A. and C.M. performed research; P.D., A.V., X.C., J.A., C.J, and C.M. analyzed data; and P.D. wrote the paper.

## Supplementary Materials

### Supplementary tables

**Table S1.** Bacterial strains used in this study

| Bacterial strains |  |  |
| --- | --- | --- |
| *L. pneumophila* CIP 108286 | [71] | Virulent *L. pneumophila* serogroup 1, strain Lens |
| *L. pneumophila* Δ*lpl0780* | [51] | Lens *lpl0780::kan* |
| *L. pneumophila* Δ*dotA* | [72] | Lens *lpl2613::kan* |
| *L. pneumophila* KS79 | [69] | JR32 derived strain |
| *L. pneumophila* KS79 Δ*icmS* | This study | JR32, *icmS::kan* |
| *L. pneumophila* CIP 107629 | [71] | Virulent *L. pneumophila* serogroup 1, strain Paris |
| *L. pneumophila* Δ*lpp0809* | This study | Paris, scar-free deletion of *lpp0809* |
| Cell line |  |  |
| U937 (ATCC number: CRL-1593.2<sup>TM</sup>) | [73] | Human monocytes |

**Table S2.** Plasmids used in this study

| Plasmids |  |  |
| --- | --- | --- |
| pXDC61 | [69] | *Legionella* expression vector carrying β-lactamase (TEM-1) gene for creation of translational gene fusions |
| pTEM-*lepA* | [19] | pXDC61 carrying β-lactamase TEM gene fused to *lepA* |
| pTEM-*fabI* | [19] | pXDC61 carrying β-lactamase TEM gene fused to *fabI* |
| pTEM-*ralF* | [19] | pXDC61 carrying β-lactamase TEM gene fused to *ralF* |
| pTEM-*legK2* | [74] | pXDC61 carrying β-lactamase TEM gene fused to *legK2* |
| pTEM-*legK4* | [74] | pXDC61 carrying β-lactamase TEM gene fused to *legK4* |
| pTEM-*lepB* | [51] | pXDC61 carrying β-lactamase TEM gene fused to *lepB* |
| pTEM-*sidM* | [51] | pXDC61 carrying β-lactamase TEM gene fused to *sidM* |
| pTEM-*sidJ* | [51] | pXDC61 carrying β-lactamase TEM gene fused to *sidJ* |
| pTEM-*lidA* | [51] | pXDC61 carrying β-lactamase TEM gene fused to *lidA* |
| pTEM-*ankX* | This study | pXDC61 carrying β-lactamase TEM gene fused to *ankX* |
| pTEM-*sidD* | This study | pXDC61 carrying β-lactamase TEM gene fused to *sidD* |
| pTEM-*ankJ* | This study | pXDC61 carrying β-lactamase TEM gene fused to *ankJ* |
| pTEM-*sidK* | This study | pXDC61 carrying β-lactamase TEM gene fused to *sidK* |
| pTEM-*ceg14* | This study | pXDC61 carrying β-lactamase TEM gene fused to *ceg14* |
| pTEM-*mavC* | This study | pXDC61 carrying β-lactamase TEM gene fused to *mavC* |
| pTEM-*mvca* | This study | pXDC61 carrying β-lactamase TEM gene fused to *mvca* |
| pHA-MCS-*mCherry* | This study | pXDC61 derivative with *blaM* gene replaced by 4 HA tags and the insertion of *mCherry* gene in operon behind the multiple cloning site |
| pHA | This study | pXDC61 derivative with *blaM* gene replaced by 4 HA tags |
| pHA-*sidM*-*mCherry* | This study | pHA-MCS-*mCherry* carrying 4HA tag fused to *sidM* |
| pHA-*lpp0809* | This study | pHA carrying 4 HA tag fused to *lpp0809* |
| pHA-*icdH* | This study | pHA carrying 4 HA tag fused to *icdH* |
| pJCL2968 | [51] | pML5PKAN |
| pJA41 | This study | pJCL2968 with *chl<sup>R</sup>* gene replaced by a *gentamycin<sup>R</sup>* gene |
| p*lpl0780* | This study | pJA41 carrying *lpl0780* gene |
| p*lpl0780*GGDEF* | This study | pJA41 carrying *lpl0780* with mutated GGDEF domain of Lpl0780 (E438D, F439I) |
| p*lpl0220* | This study | pJA41 carrying *lpl0220* gene |
| p*lpl1118* | This study | pJA41 carrying *lpl1118* gene |
| pML5PKAN-*lpp0809* | This study | pML5PKAN carrying *lpp0809* gene |
| pML5PKAN-*sfgfp* | This study | pML5PKAN carrying *sfgfp* gene |
| pM5LPKAN-4xHA-*lpp0809* | This study | pML5PKAN carrying 4 HA tag fused to *lpp0809* |
| pAA2 | This study | pXDC61 derivative with *blaM* gene removed and the insertion of *sfgfp* gene in operon behind the multiple cloning site |
| pAA2-*dotN* | This study | pAA2 carrying *dotN* gene fused to *sfgfp* gene |
| pGEM-*mazF-kan* | [75] | *E. coli* vector carrying the *mazF-kan* cassette |

**Table S3.**
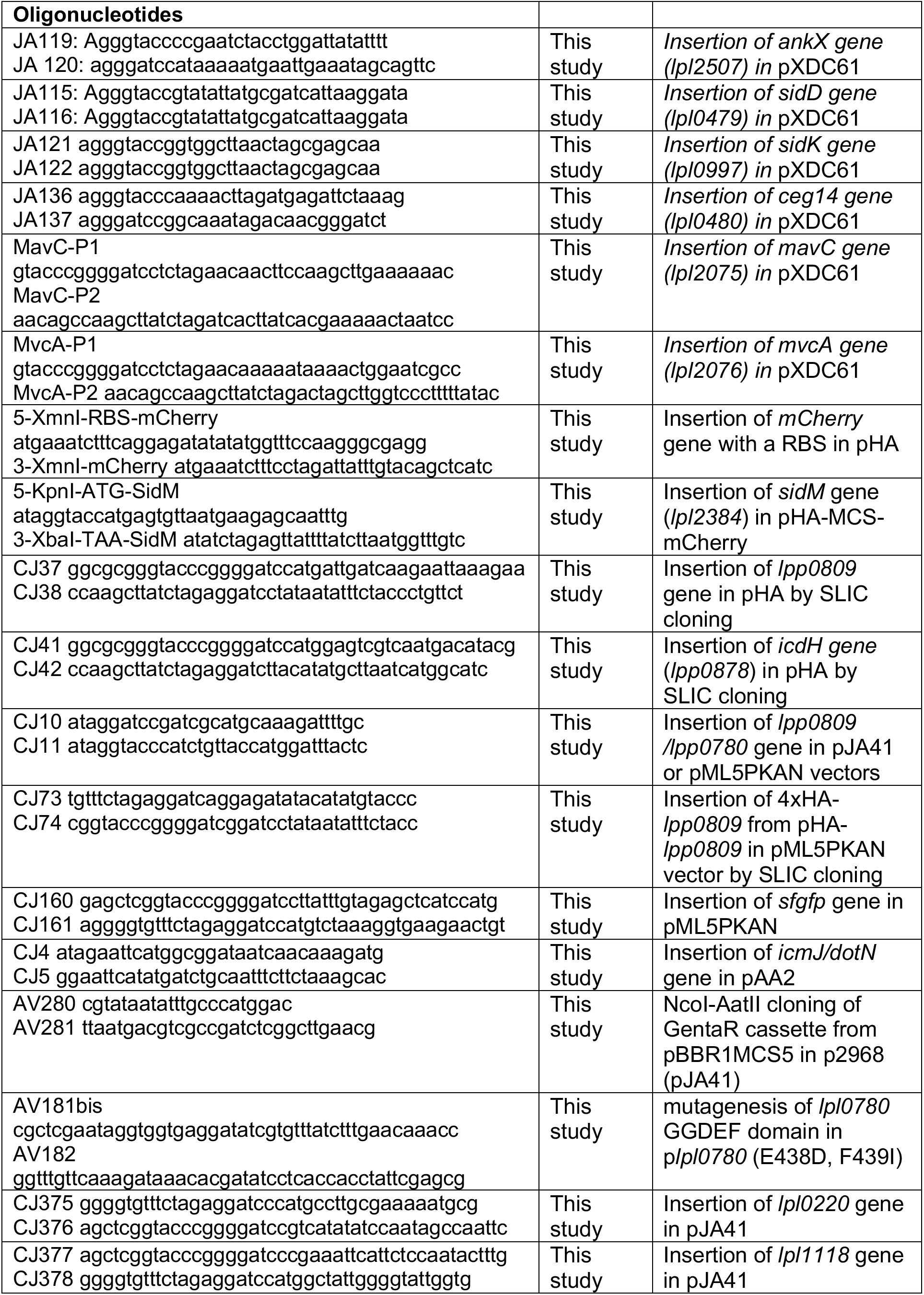
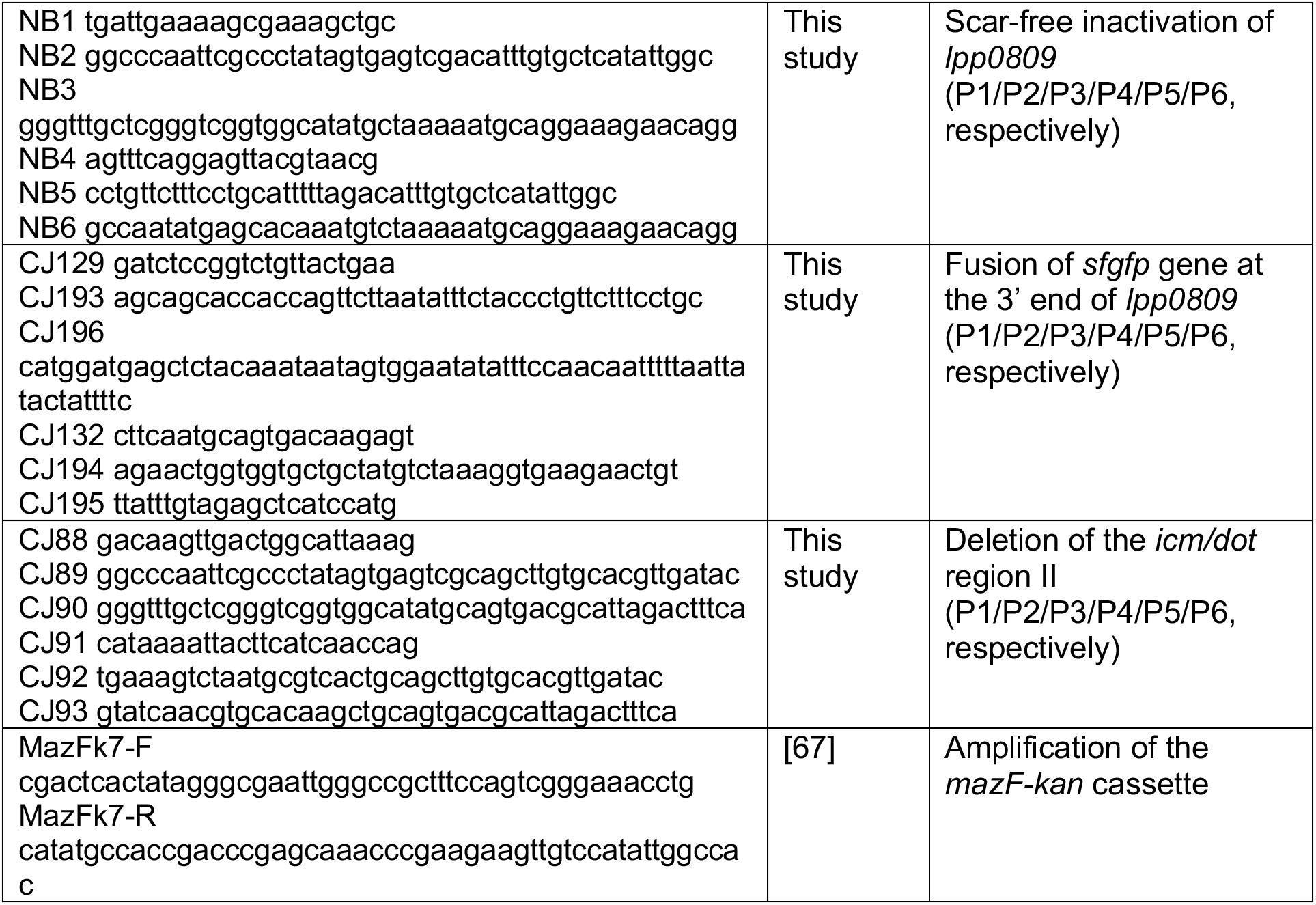
Oligonucleotides used in this study

**Table S4.** Differentially expressed genes in the *Δlpl0780* strain compared to the WT Lens strain. RNA-seq analysis was performed on three biological replicates of rRNA-depleted RNA preparations, for each strain grown on LGM medium until stationary phase (MGX-Montpellier GenomiX). RNA-seq data have been deposited in European Nucleotide Archive database at EMBL-EBI (https://www.ebi.ac.uk/ena) under accession number PRJEB33700. Alignment of RNAseq reads was performed against *Legionella pneumophila* Lens genome (NC_006369.1) using the ELAND program (CASAVA 1.8.2) and taking into account only uniquely mapped read pairs. *Differentially expressed genes were determined using DESeq2.

| Gene ID | Description | log2FoldChange* | pvalue* |
| --- | --- | --- | --- |
| *lpl2859* | hypothetical protein (hemimethylated DNA-binding protein) | -0.879520996303052 | 6.97731565361582e-07 |
| *lpl0199* | hypothetical protein | -0.764503549407963 | 8.32860031769141e-06 |
| *lpl2438* | Major facilitator family transporter | -0.69353999725098 | 2.82205150832313e-05 |
| *lpl0060* | hypothetical protein (glutaredoxin) | 0.792662085251629 | 2.33394227773472e-05 |
| *lpls01* | miscRNA | 0.75035742748646 | 2.65515318752775e-05 |
| *lpl2327* | hypothetical protein | 0.686373222423833 | 5.53639957005615e-05 |
| *lpl0980* | YchK TldD | 0.67690019830226 | 0.000121594136691985 |
| *lpl0069* | hypothetical protein | 0.62619907136437 | 0.000123752606669535 |
| *lpl0474* | hypothetical protein | -0.711649207237193 | 0.000136334248356491 |
| *lpl1801* | hypothetical protein | 0.561338139793686 | 0.00021579202365371 |
| *lpl2450* | hypothetical protein | -0.709073439548234 | 0.000193209120218638 |
| *lpl1223* | hemF (coproporphyrinogen III oxidase) | 0.733387224021937 | 0.000224986231470102 |

### Supplementary Figures

**Fig. S1.**
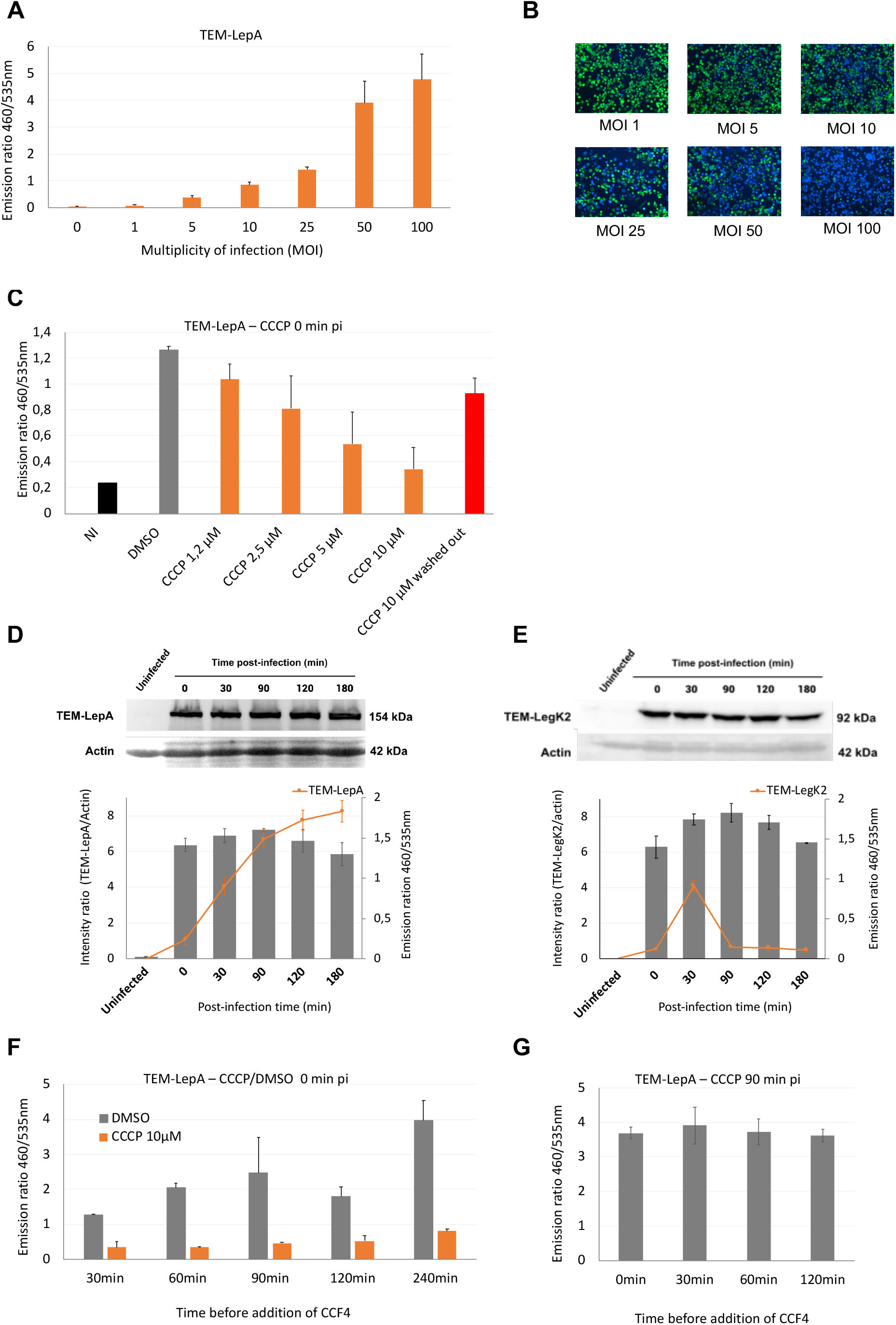
The protonophore CCCP inhibits Icm/Dot secretion activity for extended time. (A and B) Appropriate Multiplicity Of Infection (MOI)=20 for the ß-lactamase translocation assay. TEM-LepA-expressing *L. pneumophila* were used to infect U937 cells for one hour at 37°C at different MOI. Cells were then incubated with the ß-lactamase substrate CCF4/AM for 1.5 hour at room temperature. (A) Measurement of TEM-LepA translocation in U937 cells as function of MOI. (B) Fluorescence images captured at 460 and 530 nm and merged (x10). (C) Effect of CCCP or the DMSO carrier on translocation of the TEM-LepA effector in U937 cells. Different concentrations of CCCP were added at the beginning of the infection of U937 cells by TEM-LepA-producing bacteria (MOI=20). After 30 min of incubation at 37°C, cells were incubated with the ß-lactamase substrate CCF4/AM for 1.5 hour at room temperature. (D, E) Constitutive production of TEM-LepA (D) or TEM-LegK2 (E) during infection. Immunodetection of TEM in protein extract collected at different time during infection of U937 cells by TEM-LepA or TEM-LegK2-producing bacteria (MOI=20). The host protein actin was detected as a loading control (top panel). The intensity of protein bands from two independent experiments was measured by Image J and the ratios of TEM-effector to actin were plotted for each time point. The orange line represents the emission ratio of TEM-LepA or TEM-LegK2 in translocation assay from two independent experiments at each time point (bottom panel). (F) Time course effect of the CCCP protonophore or the DMSO carrier on translocation of the TEM-LepA effector in U937 cells. 10 µM CCCP or DMSO only were added at the beginning of the infection of U937 cells by TEM-LepA-producing bacteria (MOI=20). After different indicated incubation times at 37°C, cells were then incubated with the ß-lactamase substrate CCF4/AM for 1.5 hour at room temperature. (G) Stability of TEM-LepA activity in U937 cells. 10 µM CCCP were added at 1.5 hour of the infection of U937 cells by TEM-LepA-producing bacteria (MOI=20). After different indicated incubation times at 37°C, cells were then incubated with the ß-lactamase substrate CCF4/AM for 1.5 hour at room temperature. All these experiments are representative of at least 2 independent experiments.

**Fig. S2.**
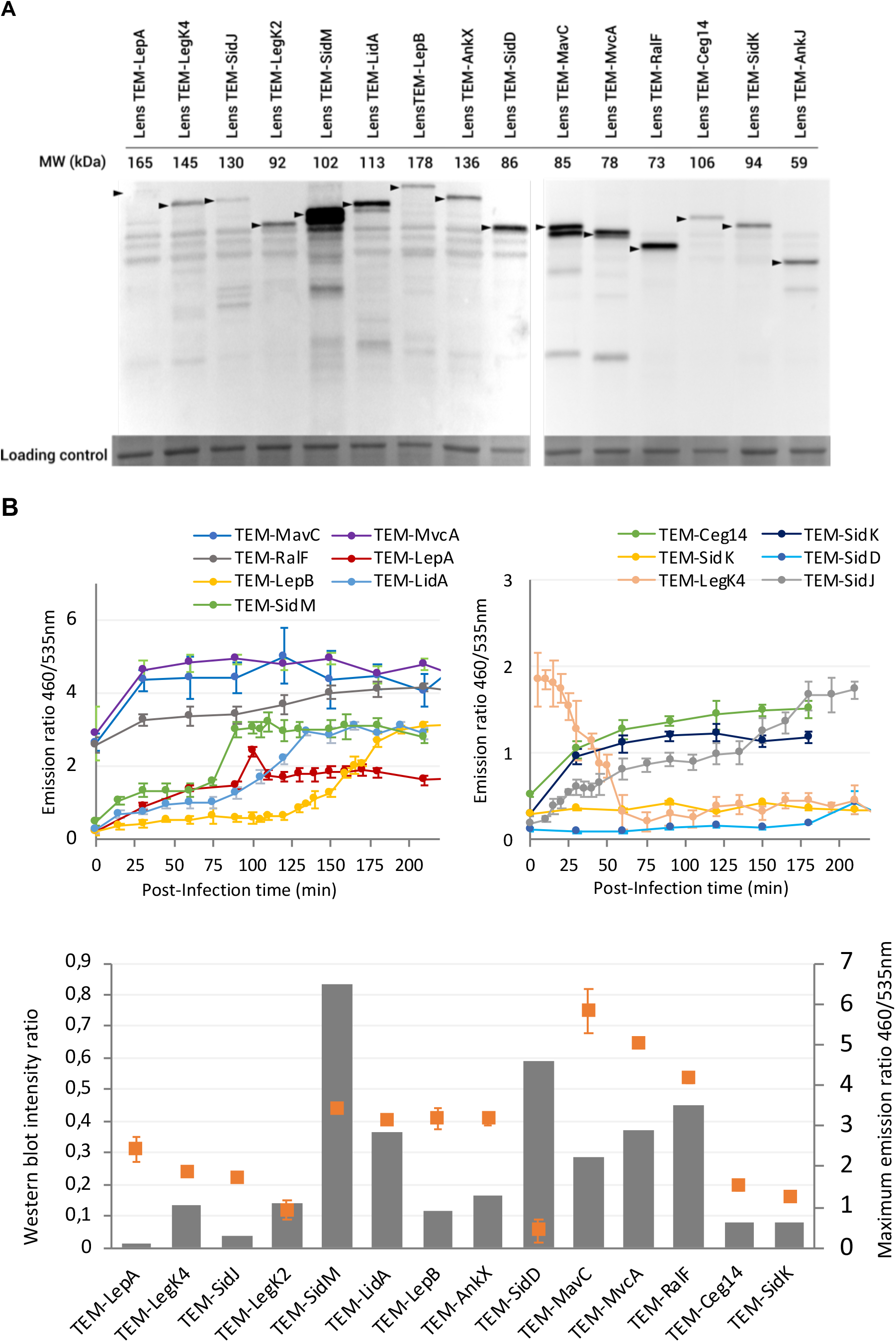
Translocation kinetic profiles are independent of the effector synthesis level. (A) The IPTG-induced TEM-effectors genes express stable protein fusions in *L. pneumophila.* Immunoblots for each TEM-effector fusion used in this study, expressed in WT pXDC61 derivative-containing Lens strains grown in the presence of 500 µM IPTG. (B) Translocation kinetic profiles of corresponding TEM-effector fusion protein measured by CCCP-combined ß-lactamase assays (top panel). The intensity of protein bands from Fig. S2A was measured by Image J and the ratios of TEM-effector to the loading control were plotted. The orange dots represent the maxima of fluorescence emission ratio (460/535nm) observed in each TEM-effector translocation assay (bottom panel). These experiments are representative of 3 independent experiments.

**Fig. S3.**
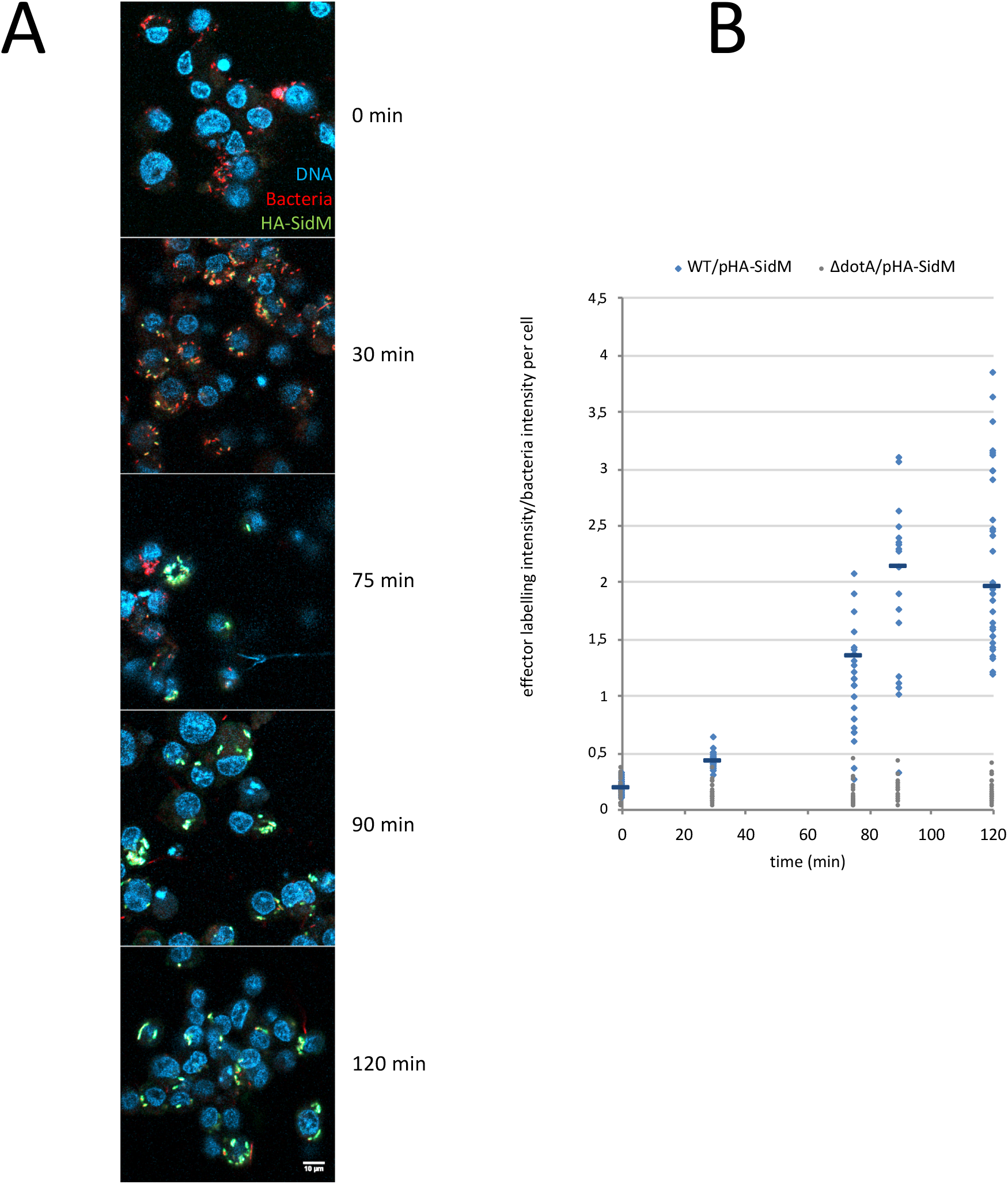
Immunodetection of SidM on LCV by immunofluorescence. (A) Confocal laser scanning micrographs of infected U937 cells after 0, 30, 75, 90 or 120 min with HA-SidM producing *L. pneumophila* (MOI 50). Nucleus were stained using DAPI (blue), bacteria expressed the fluorescent protein mCherry (red) and HA-SidM were immunolabeled with anti-HA antibodies (green). (B) Quantitative analysis of HA-SidM presence on LCV. The average intensity of HA immunolabeling was reported to the average intensity of bacteria fluorescence for each U937 cell on at least 20 cells per condition. The bar represents the median for each condition.

**Fig. S4.**
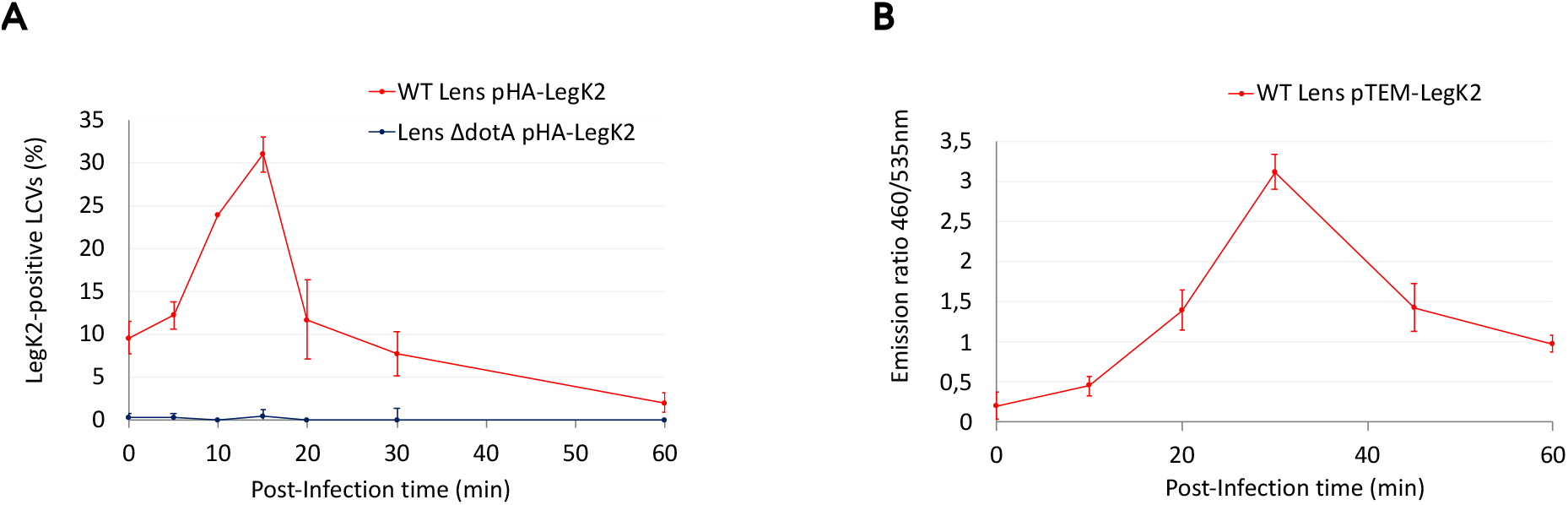
Consistency between immunodetection on LCV by immunofluorescence and secretion kinetic profile of LegK2. (A) Quantitative analysis of HA-LegK2 presence on LCV during U937 cells infection, data previously published [76]. (B) Translocation kinetic profile of TEM-LegK2 according to the CCCP-combined *β*-lactamase assay. All these experiments are representative of 2 independent experiments.

**Fig. S5.**
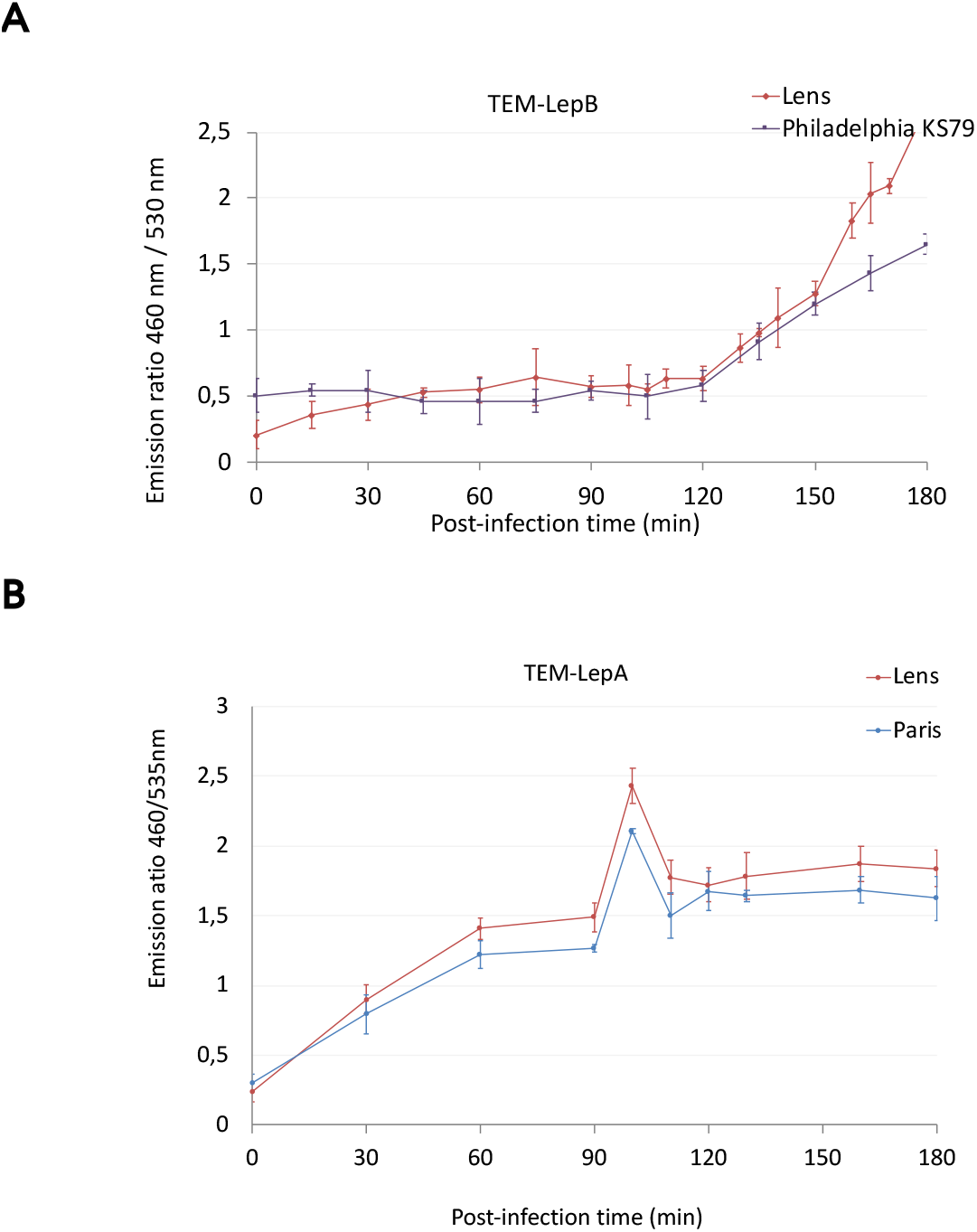
Translocation kinetic profiles are similar in different *L. pneumophila* strains. (A) Translocation kinetics of TEM-LepB in *L. pneumophila* Lens and *L. pneumophila* Philadelphia KS79 strains. (B) Translocation kinetics of TEM-LepA in *L. pneumophila* Lens and *L. pneumophila* Paris strains. These results are representative of 2 independent experiments.

**Fig. S6.**
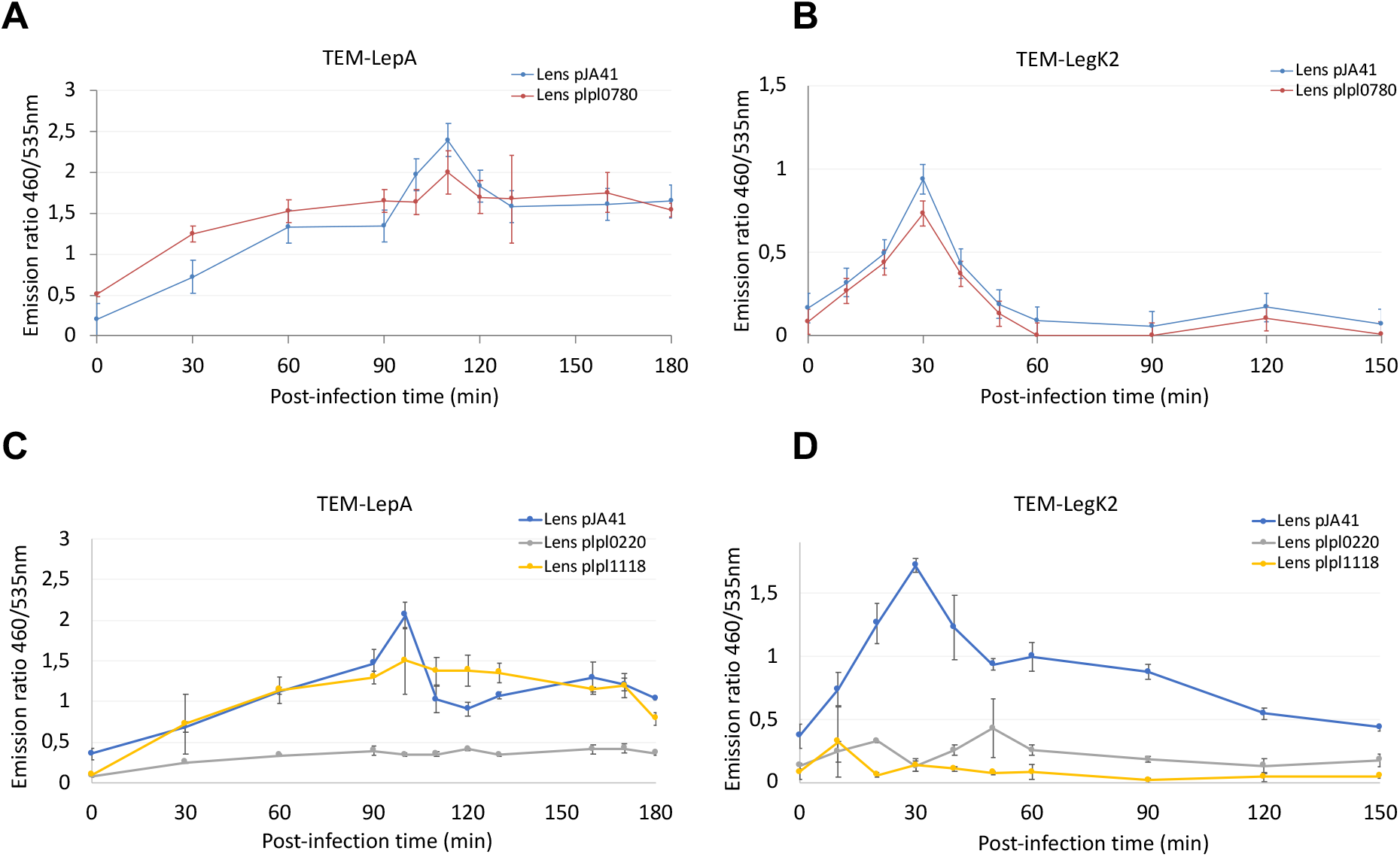
(AB) Overproduction of the DGC Lpl0780 does not alter the translocation kinetic profiles of TEM-LepA and TEM-LegK2. Translocation kinetics of TEM-LepA (A) and TEM-LegK2 (B) in *L. pneumophila* Lens containing empty plasmid pJA41 or the *lpl0780* gene expressing plasmid p*lpl0780*. These results are representative of 2 independent experiments. (CD) Overproduction of the PDE Lpl0220 and Lpl1118 inhibit the translocation of both TEM-LepA and TEM-LegK2, and the translocation of TEM-legK2, respectively. Translocation kinetics of TEM-LepA (C) and TEM-LegK2 (D) in *L. pneumophila* Lens containing empty plasmid pJA41, or the *lpl0220* gene expressing plasmid p*lpl0220,* or the *lpl1118* gene expressing plasmid p*lpl1118*. These results are representative of 2 independent experiments.

**Fig. S7.**
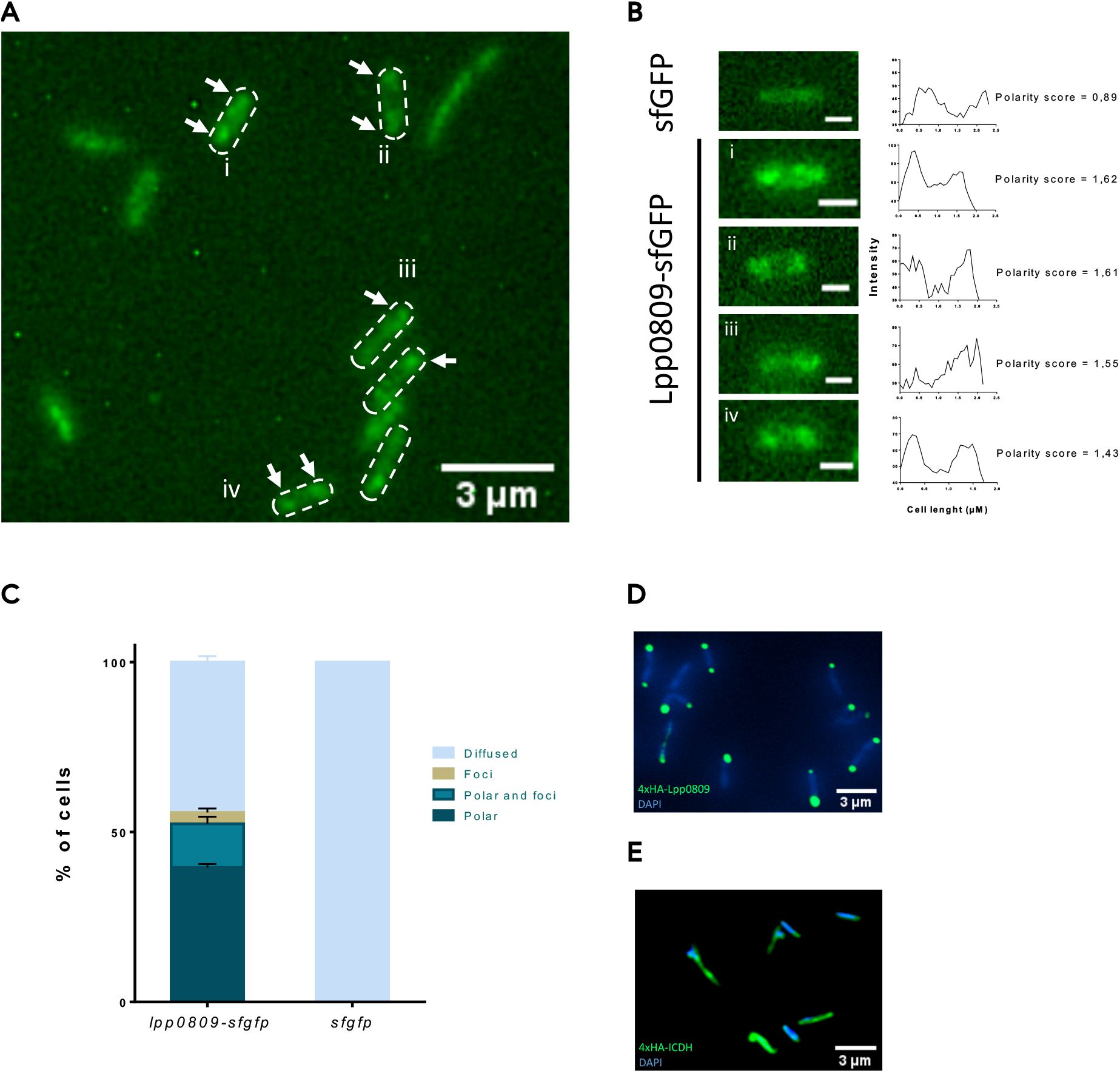
Lpp0809-sfGFP and 4xHA-tagged-Lpp0809 localize at the bacterial poles. (A) Localization of the recombinant Lpp0809-sfGFP in Paris strain expressing a chromosomal copy of *lpp0809-sfgfp* fusion (scale bar = 3µm). (B) ImageJ analysis of fluorescence intensity along the axis of representative cells (scale bar = 0,5µM). (C) Percentage of stationary phase *L. pneumophila* cells displaying cytoplasmic, foci, or polar localization of Lpp0809-sfGFP are indicated (n ≥ 200 bacteria, error bars represent the SD). (D and E) Immunodetection of a 4xHA-tagged-Lpp0809 (D) and a 4xHA-tagged ICDH (isocitrate dehydrogenase) as cytoplasmic protein control (E) by an anti-HA antibody (green) and DAPI staining (blue) in Paris strain (scale bar = 3µm). All these experiments are representative of 2 independent experiments.

**Fig. S8.**
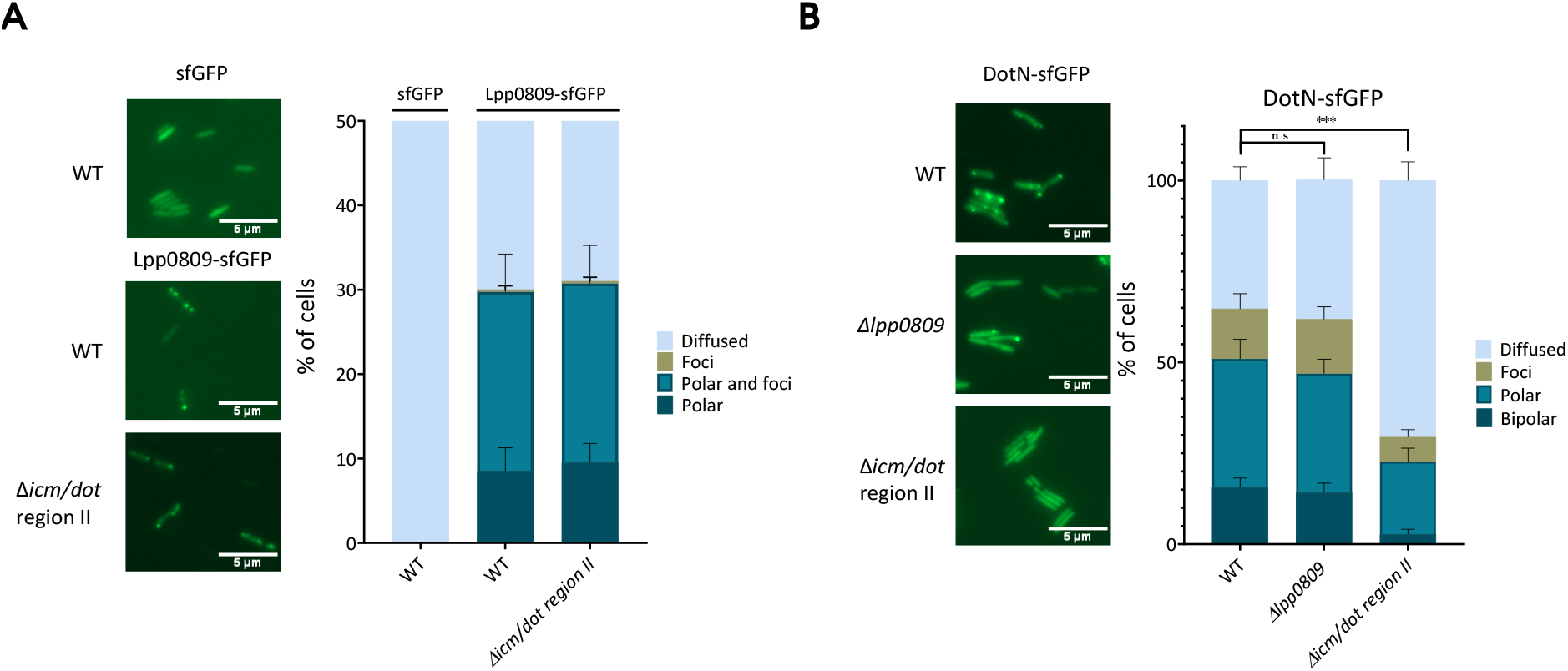
The polar localizations of Lpp0809-sfGFP and DotN-sfGFP are independent of each other. (A) Localization of the Lpp0809-sfGFP fusion protein in WT Paris strain and in Δ*icm/dot* (region II) derivative mutant strain (n ≥ 200 bacteria, error bars represent the SD). (B) Localization of the DotN-sfGFP fusion protein in WT Paris strain, Δ*lpp0809* and Δ*icm/dot* (region II) derivative mutant strains (n ≥ 200 bacteria, error bars represent the SD). All these experiments are representative of 2 independent experiments.

## References

[1] Agbor TA, McCormick BA. Salmonella effectors: important players modulating host cell function during infection. Cell Microbiol. 2011;13:1858–69.

[2] Lara-Tejero M, Kato J, Wagner S, Liu X, Galán JE. A sorting platform determines the order of protein secretion in bacterial type III systems. Science. 2011;331:1188–91.

[3] Little DJ, Coombes BK. Molecular basis for CesT recognition of type III secretion effectors in enteropathogenic Escherichia coli. PLoS Pathog. 2018;14:e1007224.

[4] Zhu W, Banga S, Tan Y, Zheng C, Stephenson R, Gately J, et al. Comprehensive identification of protein substrates of the Dot/Icm type IV transporter of Legionella pneumophila. PLoS One. 2011;6:e17638.

[5] Berger K, Isberg R. Two distinct defects in intracellular growth complemented by a single genetic locus in Legionella pneumophila. Mol Microbiol. 1993;7:7–19.

[6] Marra A, Blander SJ, Horwitz MA, Shuman HA. Identification of a Legionella pneumophila locus required for intracellular multiplication in human macrophages. Proc Natl Acad Sci U S A. 1992;89:9607–11.

[7] Vincent CD, Friedman JR, Jeong KC, Buford EC, Miller JL, Vogel JP. Identification of the core transmembrane complex of the Legionella Dot/Icm type IV secretion system. Mol Microbiol. 2006;62:1278–91.

[8] Vincent CD, Friedman JR, Jeong KC, Sutherland MC, Vogel JP. Identification of the DotL coupling protein subcomplex of the Legionella Dot/Icm type IV secretion system. Mol Microbiol. 2012;85:378–91.

[9] Kwak MJ, Kim JD, Kim H, Kim C, Bowman JW, Kim S, et al. Architecture of the type IV coupling protein complex of Legionella pneumophila. Nat Microbiol. 2017;2:17114.

[10] Meir A, Macé K, Lukoyanova N, Chetrit D, Hospenthal MK, Redzej A, et al. Mechanism of effector capture and delivery by the type IV secretion system from Legionella pneumophila. Nat Commun. 2020;11:2864.

[11] Vincent CD, Vogel JP. The Legionella pneumophila IcmS-LvgA protein complex is important for Dot/Icm-dependent intracellular growth. Mol Microbiol. 2006;61:596–613.

[12] Sutherland MC, Nguyen TL, Tseng V, Vogel JP. The Legionella IcmSW complex directly interacts with DotL to mediate translocation of adaptor-dependent substrates. PLoS Pathog. 2012;8:e1002910.

[13] Cambronne ED, Roy CR. The Legionella pneumophila IcmSW complex interacts with multiple Dot/Icm effectors to facilitate type IV translocation. PLoS Pathog. 2007;3:e188.

[14] Ghosal D, Chang YW, Jeong KC, Vogel JP, Jensen GJ. In situ structure of the Legionella Dot/Icm type IV secretion system by electron cryotomography. EMBO Rep. 2017.

[15] Chetrit D, Hu B, Christie PJ, Roy CR, Liu J. A unique cytoplasmic ATPase complex defines the Legionella pneumophila type IV secretion channel. Nat Microbiol. 2018;3:678–86.

[16] Ghosal D, Jeong KC, Chang YW, Gyore J, Teng L, Gardner A, et al. Molecular architecture, polar targeting and biogenesis of the Legionella Dot/Icm T4SS. Nat Microbiol. 2019;4:1173–82.

[17] Jeong KC, Ghosal D, Chang YW, Jensen GJ, Vogel JP. Polar delivery of Legionella type IV secretion system substrates is essential for virulence. Proc Natl Acad Sci U S A. 2017.

[18] Durie CL, Sheedlo MJ, Chung JM, Byrne BG, Su M, Knight T, et al. Structural analysis of the. Elife. 2020;9.

[19] Charpentier X, Gabay JE, Reyes M, Zhu JW, Weiss A, Shuman HA. Chemical genetics reveals bacterial and host cell functions critical for type IV effector translocation by Legionella pneumophila. PLoS Pathog. 2009;5:e1000501.

[20] Brüggemann H, Hagman A, Jules M, Sismeiro O, Dillies M, Gouyette C, et al. Virulence strategies for infecting phagocytes deduced from the in vivo transcriptional program of Legionella pneumophila. Cell Microbiol. 2006;8:1228–40.

[21] Liu Y, Gao P, Banga S, Luo ZQ. An in vivo gene deletion system for determining temporal requirement of bacterial virulence factors. Proc Natl Acad Sci U S A. 2008;105:9385–90.

[22] Ingmundson A, Delprato A, Lambright DG, Roy CR. Legionella pneumophila proteins that regulate Rab1 membrane cycling. Nature. 2007;450:365–9.

[23] Neunuebel MR, Chen Y, Gaspar AH, Backlund PS, Yergey A, Machner MP. De-AMPylation of the small GTPase Rab1 by the pathogen Legionella pneumophila. Science. 2011;333:453–6.

[24] Bardill JP, Miller JL, Vogel JP. IcmS-dependent translocation of SdeA into macrophages by the Legionella pneumophila type IV secretion system. Mol Microbiol. 2005;56:90–103.

[25] Black MH, Osinski A, Gradowski M, Servage KA, Pawłowski K, Tomchick DR, et al. Bacterial pseudokinase catalyzes protein polyglutamylation to inhibit the SidE-family ubiquitin ligases. Science. 2019;364:787–92.

[26] Gan N, Zhen X, Liu Y, Xu X, He C, Qiu J, et al. Regulation of phosphoribosyl ubiquitination by a calmodulin-dependent glutamylase. Nature. 2019;572:387–91.

[27] Bhogaraju S, Bonn F, Mukherjee R, Adams M, Pfleiderer MM, Galej WP, et al. Inhibition of bacterial ubiquitin ligases by SidJ-calmodulin catalysed glutamylation. Nature. 2019;572:382–6.

[28] Sulpizio A, Minelli ME, Wan M, Burrowes PD, Wu X, Sanford EJ, et al. Protein polyglutamylation catalyzed by the bacterial calmodulin-dependent pseudokinase SidJ. Elife. 2019;8.

[29] Kubori T, Shinzawa N, Kanuka H, Nagai H. Legionella metaeffector exploits host proteasome to temporally regulate cognate effector. PLoS Pathog. 2010;6:e1001216.

[30] Urbanus ML, Quaile AT, Stogios PJ, Morar M, Rao C, Di Leo R, et al. Diverse mechanisms of metaeffector activity in an intracellular bacterial pathogen, Legionella pneumophila. Mol Syst Biol. 2016;12:893.

[31] Aurass P, Gerlach T, Becher D, Voigt B, Karste S, Bernhardt J, et al. Life Stage-specific Proteomes of Legionella pneumophila Reveal a Highly Differential Abundance of Virulence-associated Dot/Icm effectors. Mol Cell Proteomics. 2016;15:177–200.

[32] Gan N, Guan H, Huang Y, Yu T, Fu J, Nakayasu ES, et al. Legionella pneumophila regulates the activity of UBE2N by deamidase-mediated deubiquitination. EMBO J. 2020;39:e102806.

[33] Nagai H, Cambronne ED, Kagan JC, Amor JC, Kahn RA, Roy CR. A C-terminal translocation signal required for Dot/Icm-dependent delivery of the Legionella RalF protein to host cells. Proc Natl Acad Sci U S A. 2005;102:826–31.

[34] Lifshitz Z, Burstein D, Peeri M, Zusman T, Schwartz K, Shuman HA, et al. Computational modeling and experimental validation of the Legionella and Coxiella virulence-related type-IVB secretion signal. Proc Natl Acad Sci U S A. 2013;110:E707–15.

[35] Meir A, Chetrit D, Liu L, Roy CR, Waksman G. Legionella DotM structure reveals a role in effector recruiting to the Type 4B secretion system. Nat Commun. 2018;9:507.

[36] Kim H, Kubori T, Yamazaki K, Kwak MJ, Park SY, Nagai H, et al. Structural basis for effector protein recognition by the Dot/Icm Type IVB coupling protein complex. Nat Commun. 2020;11:2623.

[37] Meir A, Mace K, Lukoyanova N, Chetrit D, Hospenthal MK, Redzej A, et al. Mechanism of effector capture and delivery by the type IV secretion system from Legionella pneumophila. Nat Commun. 2020;11:2864.

[38] Xu J, Xu D, Wan M, Yin L, Wang X, Wu L, et al. Structural insights into the roles of the IcmS-IcmW complex in the type IVb secretion system of Legionella pneumophila. Proc Natl Acad Sci U S A. 2017;114:13543–8.

[39] Jeong KC, Sutherland MC, Vogel JP. Novel export control of a Legionella Dot/Icm substrate is mediated by dual, independent signal sequences. Mol Microbiol. 2015;96:175–88.

[40] Charpentier X, Oswald E. Identification of the secretion and translocation domain of the enteropathogenic and enterohemorrhagic Escherichia coli effector Cif, using TEM-1 beta-lactamase as a new fluorescence-based reporter. J Bacteriol. 2004;186:5486–95.

[41] Qiu J, Yu K, Fei X, Liu Y, Nakayasu ES, Piehowski PD, et al. A unique deubiquitinase that deconjugates phosphoribosyl-linked protein ubiquitination. Cell Res. 2017;27:865–81.

[42] Liu Y, Luo ZQ. The Legionella pneumophila effector SidJ is required for efficient recruitment of endoplasmic reticulum proteins to the bacterial phagosome. Infect Immun. 2007;75:592–603.

[43] Machner M, Isberg R. Targeting of host Rab GTPase function by the intravacuolar pathogen Legionella pneumophila. Dev Cell. 2006;11:47–56.

[44] Murata T, Delprato A, Ingmundson A, Toomre D, Lambright D, Roy C. The Legionella pneumophila effector protein DrrA is a Rab1 guanine nucleotide-exchange factor. Nat Cell Biol. 2006;8:971–7.

[45] Müller MP, Peters H, Blümer J, Blankenfeldt W, Goody RS, Itzen A. The Legionella effector protein DrrA AMPylates the membrane traffic regulator Rab1b. Science. 2010;329:946–9.

[46] Hardiman CA, Roy CR. AMPylation is critical for Rab1 localization to vacuoles containing Legionella pneumophila. MBio. 2014;5:e01035–13.

[47] Neunuebel MR, Mohammadi S, Jarnik M, Machner MP. Legionella pneumophila LidA affects nucleotide binding and activity of the host GTPase Rab1. J Bacteriol. 2012;194:1389–400.

[48] Cheng W, Yin K, Lu D, Li B, Zhu D, Chen Y, et al. Structural insights into a unique Legionella pneumophila effector LidA recognizing both GDP and GTP bound Rab1 in their active state. PLoS Pathog. 2012;8:e1002528.

[49] Mills E, Baruch K, Charpentier X, Kobi S, Rosenshine I. Real-time analysis of effector translocation by the type III secretion system of enteropathogenic Escherichia coli. Cell Host Microbe. 2008;3:104–13.

[50] Chen J, Reyes M, Clarke M, Shuman HA. Host cell-dependent secretion and translocation of the LepA and LepB effectors of Legionella pneumophila. Cell Microbiol. 2007;9:1660–71.

[51] Allombert J, Lazzaroni JC, Baïlo N, Gilbert C, Charpentier X, Doublet P, et al. Three antagonistic cyclic di-GMP-catabolizing enzymes promote differential Dot/Icm effector delivery and intracellular survival at the early steps of Legionella pneumophila infection. Infect Immun. 2014;82:1222–33.

[52] Levi A, Folcher M, Jenal U, Shuman HA. Cyclic Diguanylate Signaling Proteins Control Intracellular Growth of Legionella pneumophila. MBio. 2011;2.

[53] Jenal U, Reinders A, Lori C. Cyclic di-GMP: second messenger extraordinaire. Nat Rev Microbiol. 2017.

[54] Kubori T, Hubber AM, Nagai H. Hijacking the host proteasome for the temporal degradation of bacterial effectors. Methods Mol Biol. 2014;1197:141–52.

[55] Kubori T, Hyakutake A, Nagai H. Legionella translocates an E3 ubiquitin ligase that has multiple U-boxes with distinct functions. Mol Microbiol. 2008;67:1307–19.

[56] Weber S, Ragaz C, Reus K, Nyfeler Y, Hilbi H. Legionella pneumophila exploits PI(4)P to anchor secreted effector proteins to the replicative vacuole. PLoS Pathog. 2006;2:e46.

[57] Ivanov SS, Charron G, Hang HC, Roy CR. Lipidation by the host prenyltransferase machinery facilitates membrane localization of Legionella pneumophila effector proteins. J Biol Chem. 2010;285:34686–98.

[58] Trampari E, Stevenson CE, Little RH, Wilhelm T, Lawson DM, Malone JG. Bacterial Rotary Export ATPases are Allosterically Regulated by the Nucleotide Second Messenger Cyclic-di-GMP. J Biol Chem. 2015.

[59] Krasteva PV, Bernal-Bayard J, Travier L, Martin FA, Kaminski PA, Karimova G, et al. Insights into the structure and assembly of a bacterial cellulose secretion system. Nat Commun. 2017;8:2065.

[60] Dahlstrom KM, Giglio KM, Sondermann H, O’Toole GA. The Inhibitory Site of a Diguanylate Cyclase Is a Necessary Element for Interaction and Signaling with an Effector Protein. J Bacteriol. 2016;198:1595–603.

[61] Dahlstrom KM, Giglio KM, Collins AJ, Sondermann H, O’Toole GA. Contribution of Physical Interactions to Signaling Specificity between a Diguanylate Cyclase and Its Effector. MBio. 2015;6:e01978–15.

[62] Whitney JC, Howell PL. Synthase-dependent exopolysaccharide secretion in Gram-negative bacteria. Trends Microbiol. 2013;21:63–72.

[63] Whitney JC, Whitfield GB, Marmont LS, Yip P, Neculai AM, Lobsanov YD, et al. Dimeric c-di-GMP is required for post-translational regulation of alginate production in Pseudomonas aeruginosa. J Biol Chem. 2015;290:12451–62.

[64] Hengge R, Häussler S, Pruteanu M, Stülke J, Tschowri N, Turgay K. Recent Advances and Current Trends in Nucleotide Second Messenger Signaling in Bacteria. J Mol Biol. 2019;431:908–27.

[65] Li MZ, Elledge SJ. SLIC: a method for sequence- and ligation-independent cloning. Methods Mol Biol. 2012;852:51–9.

[66] Ish-Horowicz D, Burke JF. Rapid and efficient cosmid cloning. Nucleic Acids Res. 1981;9:2989–98.

[67] Bailo N, Kanaan H, Kay E, Charpentier X, Doublet P, Gilbert C. Scar-Free Genome Editing in Legionella pneumophila. Methods Mol Biol. 2019;1921:93–105.

[68] Segal G, Shuman HA. Characterization of a new region required for macrophage killing by Legionella pneumophila. Infect Immun. 1997;65:5057–66.

[69] de Felipe KS, Glover RT, Charpentier X, Anderson OR, Reyes M, Pericone CD, et al. Legionella eukaryotic-like type IV substrates interfere with organelle trafficking. PLoS Pathog. 2008;4:e1000117.

[70] Hiraga S, Ichinose C, Niki H, Yamazoe M. Cell cycle-dependent duplication and bidirectional migration of SeqA-associated DNA-protein complexes in E. coli. Mol Cell. 1998;1:381–7.

[71] Cazalet C, Rusniok C, Brüggemann H, Zidane N, Magnier A, Ma L, et al. Evidence in the Legionella pneumophila genome for exploitation of host cell functions and high genome plasticity. Nat Genet. 2004;36:1165–73.

[72] Ferhat M, Atlan D, Vianney A, Lazzaroni JC, Doublet P, Gilbert C. The TolC protein of Legionella pneumophila plays a major role in multi-drug resistance and the early steps of host invasion. PLoS One. 2009;4:e7732.

[73] Sundström C, Nilsson K. Establishment and characterization of a human histiocytic lymphoma cell line (U-937). Int J Cancer. 1976;17:565–77.

[74] Hervet E, Charpentier X, Vianney A, Lazzaroni JC, Gilbert C, Atlan D, et al. Protein kinase LegK2 is a type IV secretion system effector involved in endoplasmic reticulum recruitment and intracellular replication of Legionella pneumophila. Infect Immun. 2011;79:1936–50.

[75] Attaiech L, Boughammoura A, Brochier-Armanet C, Allatif O, Peillard-Fiorente F, Edwards RA, et al. Silencing of natural transformation by an RNA chaperone and a multitarget small RNA. Proc Natl Acad Sci U S A. 2016;113:8813–8.

[76] Michard C, Sperandio D, Bailo N, Pizarro-Cerda J, LeClaire L, Chadeau-Argaud E, et al. The Legionella Kinase LegK2 Targets the ARP2/3 Complex To Inhibit Actin Nucleation on Phagosomes and Allow Bacterial Evasion of the Late Endocytic Pathway. Mbio. 2015;6:14.

